# Native metabolomics identifies pteridines as CutA ligands and modulators of copper binding

**DOI:** 10.1101/2025.04.03.647050

**Authors:** Berenike C. Wagner, Christian Geibel, Amelie Stadelmann, Johanna Rapp, Hannes Link, Eva Nussbaum, Heinz-Paul Grenzendorfer, Reinhard Albrecht, Marcus D. Hartmann, Karl Forchhammer, Khaled A. Selim, Chambers C. Hughes, Daniel Petras

**Affiliations:** Organismic Interactions, Interfaculty Institute of Microbiology and Infection Medicine (IMIT), University of Tuebingen, Germany; Cluster of Excellence EXC 2124: Controlling Microbes to Fight Infection (CMFI), University of Tuebingen, Germany; Bacterial Metabolomics, Interfaculty Institute of Microbiology and Infection Medicine (IMIT), University of Tuebingen, Germany; German Center for Infection Research (DZIF), Partner Site Tuebingen, Germany; Department of Protein Evolution, Max Planck Institute for Biology Tuebingen, Germany; Microbial Biochemistry Group, Institute of Phototrophic Microbiology, Heinrich-Heine University Düsseldorf, Germany; Department of Biochemistry, University of California Riverside, USA; Interfaculty Institute of Biochemistry, University of Tuebingen, Germany; Department of Microbial Bioactive Compounds, Interfaculty Institute of Microbiology and Infection Medicine (IMIT), University of Tuebingen, Germany

**Keywords:** CutA, native metabolomics, copper, pteridines, tetrahydrobiopterin, PII-like proteins

## Abstract

CutA, a conserved protein across all domains of life, has long been linked to copper tolerance in *Escherichia coli*, though recent studies question this association. To clarify its function, we studied *cutA* knockout mutants from two phylogenetically distant species, *E. coli* and *Synechococcus elongatus* PCC 7942, using phenotyping combined with targeted and untargeted metabolomics. Native metabolomics of cell extracts revealed the lumazine dehydroxyxanthopterin B2, a previously uncharacterized pteridine, to bind CutA in both species. Based on these results, we identified other pteridines, including the essential cofactor tetrahydrobiopterin, as ligands of CutA proteins. In the presence of pterins, we observed higher affinity of CutA to copper ions. These findings, alongside the known role of pteridines as redox shuttles, suggest a previously unrecognized role for CutA in coordinating copper homeostasis and redox balance via pteridine metabolism.

**Significance:** We identified the molecular class of pteridines as natural ligands of CutA, including the so far unknown lumazine dehydroxyxanthopterin B2. Pteridines are known redox shuttles involved in various cellular processes such as cofactors for redox enzymes. Our data showed increased copper binding to CutA in the presence of pteridines. Together, these results suggest that pteridines are physiological ligands of CutA that may modulate copper binding and redox homeostasis.

## Introduction

Advances in next-generation sequencing and the application of mass spectrometry to high-throughput biological studies have revolutionized our understanding of biology. Yet, a substantial fraction of genes and proteins remain functionally uncharacterized. One of those enigmatic proteins is CutA, first identified in *Escherichia coli* as part of a gene cluster consisting of <u>c</u>opper <u>u</u>ptake and <u>t</u>ransport (*cut*) genes^1,2^. CutA’s conservation across all domains of life^3,4^, implies a crucial biological function. It is part of a widespread signal transduction superfamily of the homo-trimeric PII-like proteins^5,4,6^, with each monomer weighing approximately 12 kDa. Despite similar core structures and trimeric assembly, CutA does not share any sequence homology with the nitrogen (N)- and carbon (C)-sensing PII proteins^4,5^—key regulators of C/N homeostasis^7–10^—it lacks amino acid sequence homology, defining CutA as a PII-like protein^4,8^. In mammals, CutA is implicated in β-amyloid precursor processing via its interaction with β-site APP cleaving enzyme 1 (BACE1)^11,12^. CutA also co-purifies with acetylcholinesterase (AChE)^13^ and exhibits a similar expression pattern in the brain^14^, where it is described to facilitate AChE folding, oligomerization, secretion^15^ and surface localization^14^. Due to its conservation across diverse organisms, from ancient cyanobacteria to eukaryotes including humans, CutA proteins are proposed to fulfill a fundamental role analogous to PII proteins in sensing and regulating cellular homeostasis.

*Escherichia coli* mutants in the *cut* cluster, categorized as *cutA–cutF*, display altered copper responses. The *cutA* locus encodes three proteins, including the cytoplasmic trimeric CutA1 (CutA in the following), the membrane-associated CutA2/DsbD (which transfers electrons from the cytoplasm to the periplasm of *E. coli*, preserving cysteine thiols in their reduced state^16^) and CutA3. Deletion mutants of the *cutA* locus exhibit hypersensitivity to metals and increased copper uptake^2^, although the contribution of *cutA1* to copper tolerance remains debated^4^, with putative pleiotropic effects of the neighboring *dsbd* complicating interpretations^3^. Structurally, *E. coli* CutA is described to bind Cu(II) via a His2Cys coordination^3^, potentially mediating redox regulation of thiol groups in copper and zinc transport ATPases^3,17^. The binding site in CutA shares structural similarity with the ATP-binding site of PII proteins^3,18,19^. Arnesano et al. (2003) proposed that CutA may function as a sensor or regulator in copper homeostasis, potentially mitigating excess copper levels through direct binding or by modulating copper import or export via interactions with membrane transporters^17^. However, recent phenotyping studies indicated that CutA is not essential for copper tolerance in either *E. coli* or the cyanobacterium *Synechococcus elongatus*^4,20^. These findings, along with CutA’s conserved structure resembling signal-transducing PII proteins, its broad distribution, and potential copper interactions, motivated us to explore its biological function in greater depth.

Our research focused on two distinct bacterial taxa to identify a potential common function: *Escherichia coli* BW25113 (*E. coli*) and *Synechococcus elongatus* PCC 7942 (*S. elongatus*). By exposing both taxa to various stress conditions, we aimed to shed light on the functional roles of the CutA protein in different environmental contexts. Building on the known extreme heat stability of many CutA proteins^21–23^, we explored their response to this stress condition. Next, we used a range of metabolomics approaches to investigate potential changes in intracellular metabolomes and search for CutA binding molecules. Using native metabolomics^24,25^, we could identify a set of putative CutA ligands, which we verified using orthogonal biophysical characterization methods. Using nuclear magnetic resonance (NMR) spectroscopy and high-resolution tandem mass spectrometry (MS/MS), we solved the structures of a novel pteridine derivative, which ultimately identified pteridines as CutA ligands and putative regulators of copper affinity and cellular redox homeostasis.

## Results

### CutA deletion alters tolerance and metabolite homeostasis during heat stress

Given CutA’s exceptional thermal stability^22,26^, we investigated its role under environmental stress conditions. Exposure to 50 °C revealed that recovery of the *S. elongatus ΔcutA::kan* mutant was severely impaired compared to the WT (**Fig. 1a**). Similarly, the *ΔcutA::kan* mutant exhibited increased susceptibility to ampicillin-induced cell wall stress, with pronounced effects observed after two days of treatment (**Fig. 1b**). A similar sensitivity was observed on carbenicillin plates (**Supplementary Fig. 1**). This heightened sensitivity may be attributed to elevated levels of reactive oxygen species (ROS) generated in response to β-lactam exposure^27–29^, potentially exacerbating cellular damage in the absence of CutA-mediated protective mechanisms. Repetition of the experiments and full plates are shown in **Supplementary Fig. 2-3**. Global metabolomic profiling of *S. elongatus* revealed significant metabolic differences between the WT and *ΔcutA* strains. A principal coordinate analysis (PCoA) revealed significant differences in the observed metabolomes between the two groups (PERMANOVA, *p* = 0.017, **Fig. 1c**).

**Figure 1:**
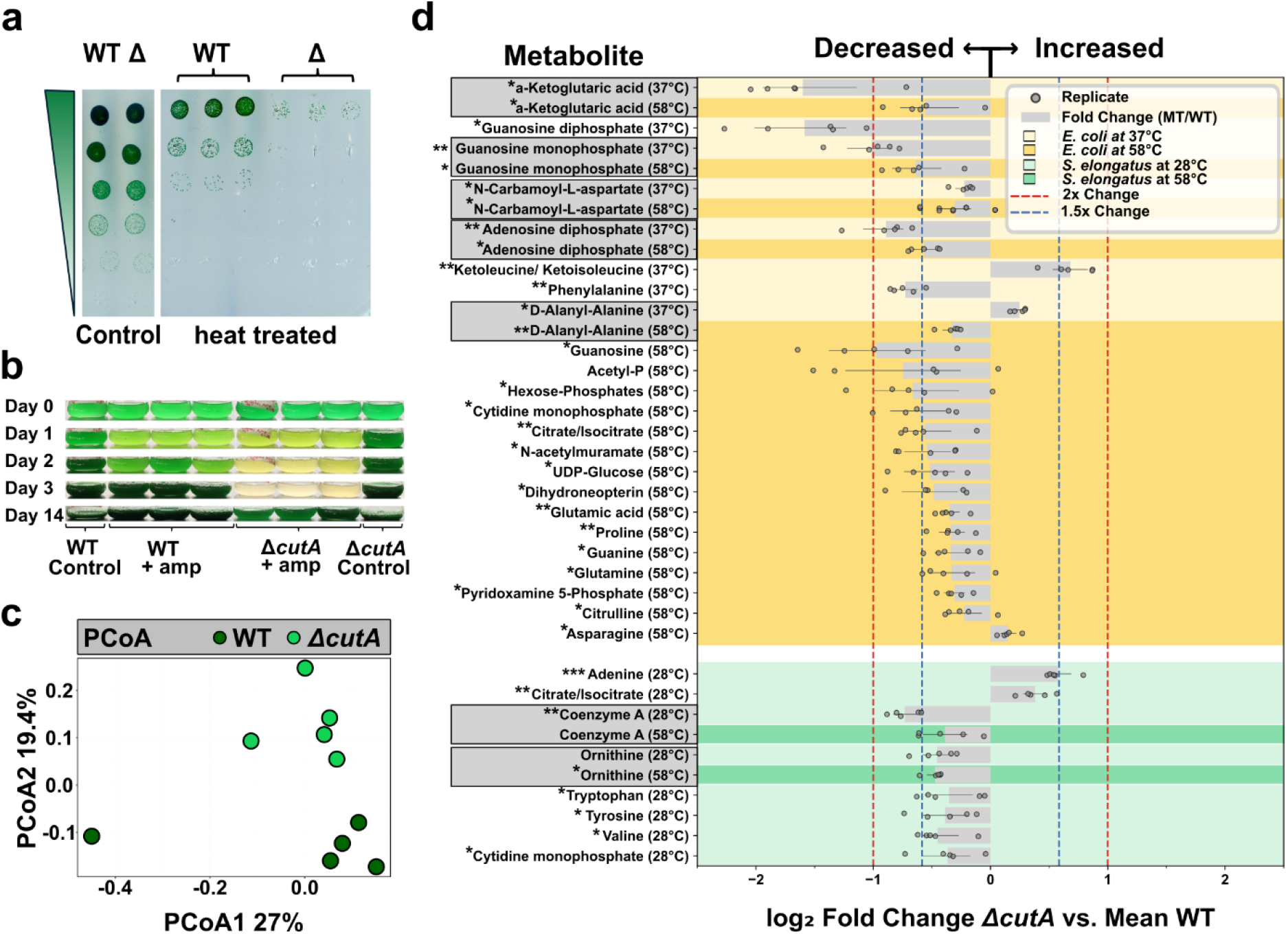
Role of CutA in heat and cell envelope stress tolerance. **a**, Viability drop dilution assay shows impaired recovery of the ΔcutA::kan mutant after heat stress. S. elongatus WT and ΔcutA::kan cultures were exposed to 50 °C for 1.5 h, serially diluted (1:10, top to bottom), and incubated at 28 °C for 7 days under constant light (30-60 µmol photons m^2^ s^-1^E), n=3 **b**, Ampicillin sensitivity of the ΔcutA::kan mutant. The growth of cultures treated with 10 µg/mL ampicillin was monitored over 14 days. Controls without treatment are shown for comparison, n=3 **c**, Principal coordinates analysis (PCoA) reveals significant divergence in the global metabolome of ΔcutA::kan compared to the WT (PERMANOVA, p = 0.017, permutations=999, R2=0.1806, PERMDISP p=0.3067). Blanks (medium controls) were subtracted before analysis, n=5 **d**, Targeted metabolomic analysis with and without heat stress. Significantly different concentrations (p-values below 0.05) of metabolites in the ΔcutA mutants normalized to the WT are shown for S. elongatus (28 °C, 58 °C) and E. coli (37 °C, 58 °C). Mean value and standard deviation of five biological replicates are shown. Each replicate was divided by the mean of the WT and shown additionally as a single dot. Metabolites of E. coli and S. elongatus are depicted in yellow and green, respectively, with brighter colors representing standard growth conditions and darker colors representing heat treatment. Grey metabolite name labels indicate significant effects observed in both treatments. Blue and red dotted lines denote thresholds for 1.5-fold and 2-fold changes, respectively. Data are presented as mean ± s.d. for five biological replicates. Statistical analysis was performed using unpaired, two-tailed t-tests. Differences between means with p-values < 0.05, < 0.01, and < 0.001 are indicated by one, two, or three stars, respectively.

Targeted metabolomic analysis of *E. coli* and *S. elongatus* under heat stress (58 °C) and control conditions revealed a downregulation of several metabolites in the *ΔcutA* mutants, particularly after heat treatment, some of which are linked to stress resistance pathways (**Fig. 1d**). Notably, dihydroneopterin and pyridoxamine 5’-phosphate, precursors for tetrahydrofolate and tetrahydrobiopterin biosynthesis, were among the affected metabolites in *E. coli* during heat treatment. This suggests impaired cofactor biosynthesis, which could compromise the cell’s ability to mitigate oxidative stress that is generated under thermal stress^30–32^, as both tetrahydrofolate and tetrahydrobiopterin are essential for maintaining redox balance and counteracting reactive oxygen species^33–36^. Together, these findings indicate that under conditions of heat or cell wall stress, the loss of *cutA* disrupts metabolite homeostasis, thereby impairing the survival of the *ΔcutA* mutants.

### Native Metabolomics uncovers CutA Binders in Cyanobacterial Cell Extracts

To further investigate the mechanisms underlying the physiological and metabolomic changes in the *ΔcutA* mutants, we utilized a native metabolomics approach that enables rapid screening of potential binding molecules from crude cell extracts (**Fig. 2a**)^24^.

**Figure 2:**
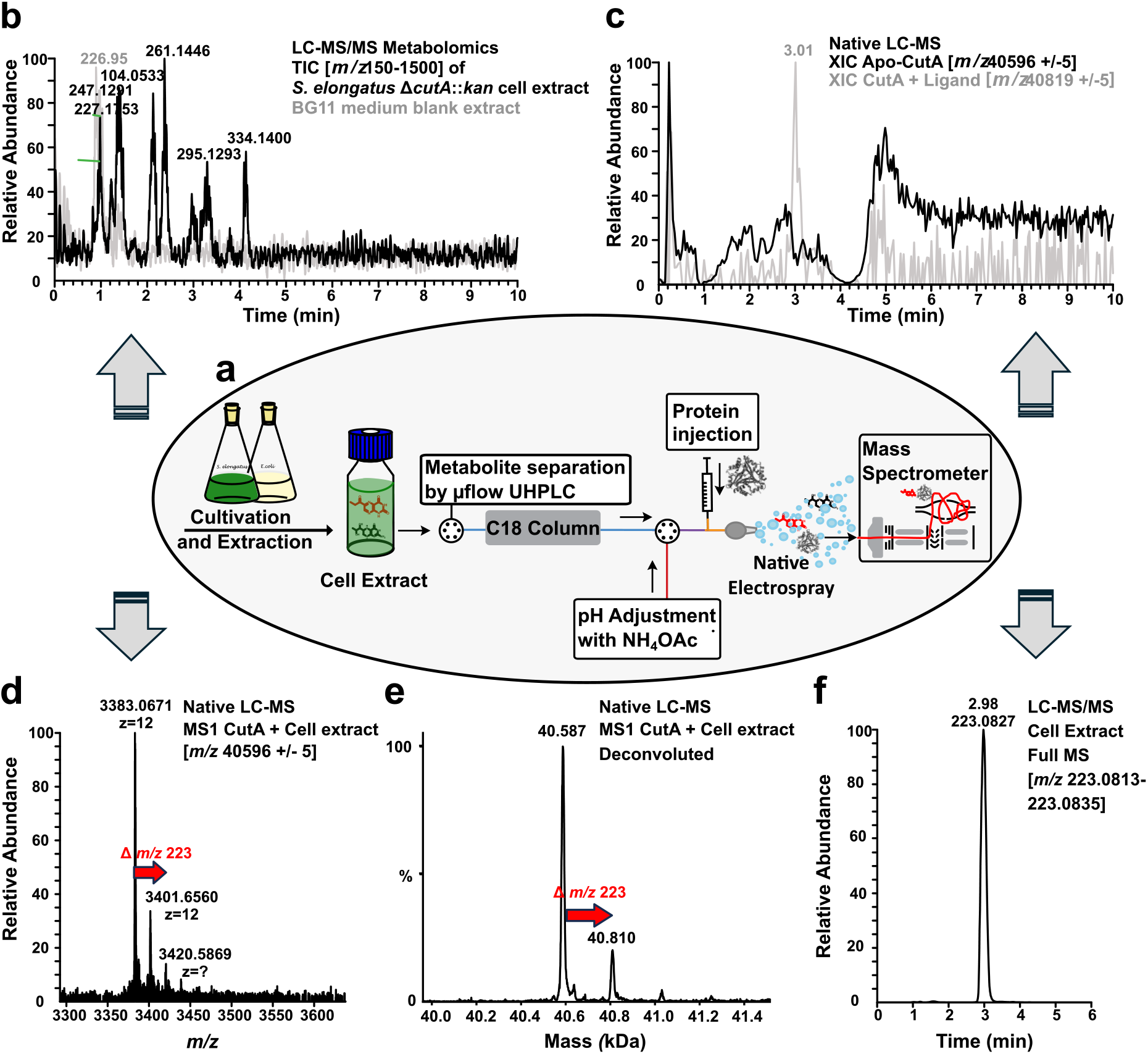
Native MS setup and identification of a putative CutA ligand using native metabolomics. **a**, E. coli CutA protein was analyzed under native conditions and screened against crude cell extracts of S. elongatus and E. coli WT or ΔcutA. Native MS Setup^24^: Cellular extracts were separated using µ-flow UHPLC. pH adjustment to near-physiological conditions (via ammonium acetate) preserved the protein’s native state. Eluting metabolites interact with the infused protein, allowing detection of protein-ligand complexes by mass spectrometry. A separate metabolomics run (high-resolution UHPLC-MS/MS) was performed without protein infusion to identify the metabolites separately. **b**, Total Ion Chromatogram (TIC) of S. elongatus cell extract (black) and BG11 medium (control, grey), revealing the metabolomic profile. **c**, Extracted Ion Chromatogram (XIC) of apo-CutA (black) and CutA bound to ligand (grey). The decrease in the XIC of apo-CutA at RT = 3.01 min (depicted in black) corresponds to an increase in the peak for the protein-ligand complex at the same retention time (depicted in grey). **d**, Mass spectrum of the E. coli CutA trimer infused to S. elongatus metabolites. In addition to the CutA trimer (3383.0671 × 12 Da, including the Strep tag), a new peak appears at 3401.6560 × 12 Da. The mass shift between the protein-ligand complex (m/z 40,819.9 ± 5) and the unbound protein (m/z 40,596.8 ± 5) suggested a molecular weight of 223 ± 2 Da for the putative binder. **e**, Deconvoluted spectrum showing the shift from apo-CutA to the protein-ligand complex. **f**, Metabolomics run of cell extract without protein infusion showing the retention time and mass corresponding to the protein-ligand binding event observed in panel 2d.

Here, crude bacterial cell extracts of *E. coli* or *S. elongatus* were injected and separated via reversed-phase chromatography. Prior to native electrospray ionization (ESI) and subsequent data acquisition, the post-column pH was adjusted to near physiological (∼ pH 6.1) conditions using ammonium acetate buffer. Metabolites eluting from the column interact with the native protein and binding can be measured as mass shifts of the apo-protein via native MS. **Figure 2b** presents the Total Ion Chromatogram (TIC) of *S. elongatus* cell extracts compared to BG11 medium extracts. The cell extracts exhibited a greater number of peaks, reflecting a higher diversity of compounds, particularly polar metabolites, eluting in the early stages of the chromatographic gradient. Over 1,000 molecular features were detected in *S. elongatus* extracts and more than 1,500 in *E. coli* extracts (**Supplementary Fig. 4**).

**Figure 2c** overlays the signal of *E. coli* apo-CutA (black), measured without metabolites, and the CutA-ligand complex (grey), observed during the analysis of CutA with metabolites from cell extracts. A marked reduction in the apo-CutA signal (black) coincides with the appearance of a prominent peak corresponding to the CutA-ligand complex (grey), indicating ligand binding. Specifically, a mass shift from 40,596.8 ± 5 Da (apo-CutA) to 40,819.9 ± 5 Da (ligand-bound CutA) was observed at a retention time (RT) of 2.98 minutes in the measurement of CutA protein with *S. elongatus* cell extract (**Fig. 2d**). The deconvoluted spectrum is shown in **Figure 2e**. A deconvoluted spectrum of *S. elongatus* CutA-ligand binding is shown in **Supplementary Fig. 5**.

The *m/z* difference was used to calculate the ligand’s molecular weight. An additional LC-MS/MS run without protein, but identical LC gradient facilitated the identification of putative ligands by comparing retention time, molecular weight, and MS/MS fragmentation patterns. In this case, a metabolite corresponding to the observed mass shift (plus a proton from ionization) was detected at 2.98 minutes in the chromatogram of the extract analyzed without protein (**Fig. 2f**).

The MS/MS spectrum of this compound is shown in **Figure 3b**. Based on its mass and retention time, the putative binder was identified as a small polar compound. MS/MS analysis of the ion (monoisotopic mass, MR = 222.0753 Da; *m/z* = 223.0824 [M+H]^+^) revealed a molecular formula of C_9_H_10_N_4_O_3_. Subsequent database searches yielded over ten potential structures; however, the idea that this binder represented a new chemical structure was also a distinct possibility.

**Figure 3:**
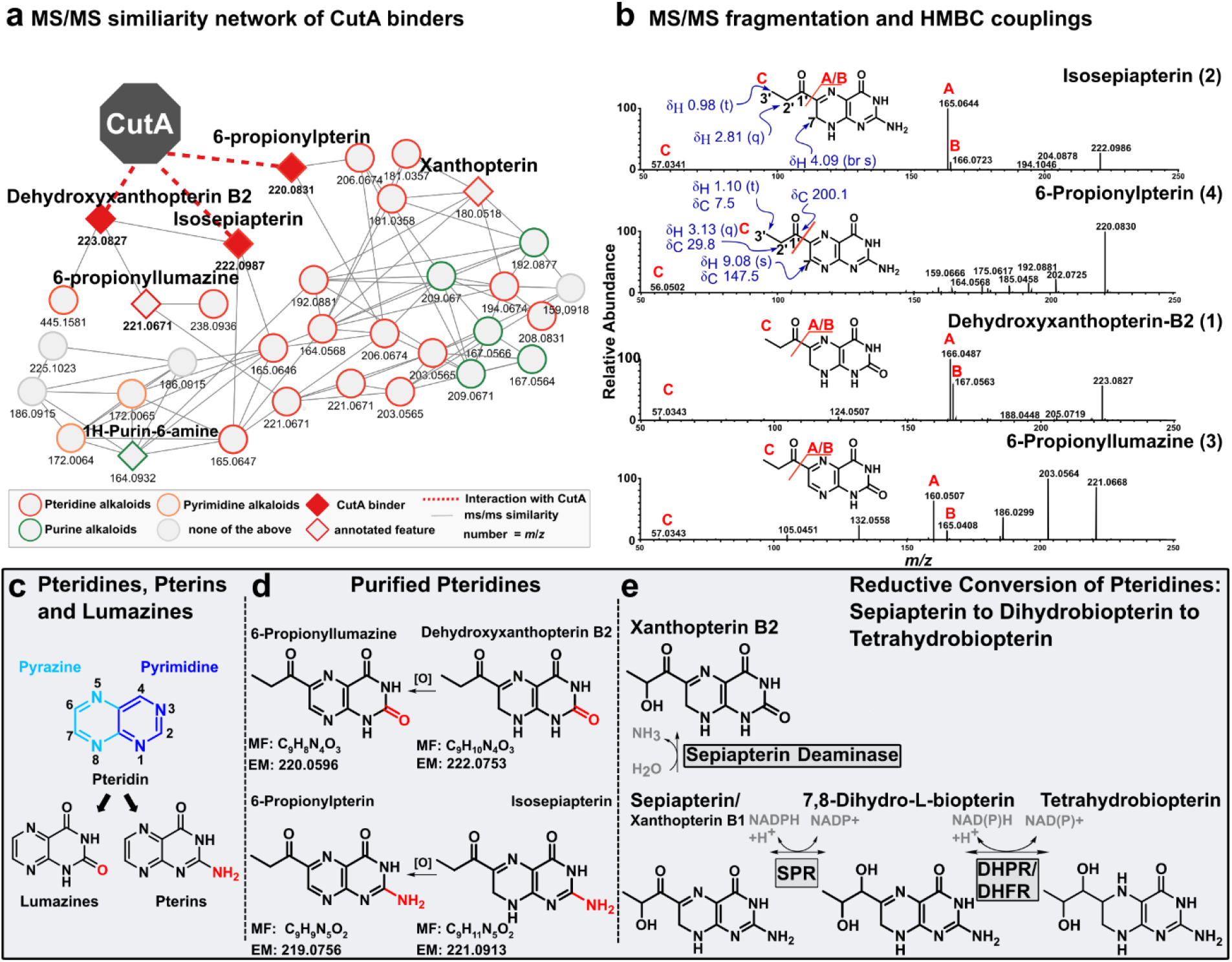
Characterization and Structural Analysis of Putative CutA binder and Co-purified Pteridines. **a**, Molecular correlation network based on MS/MS similarity for HPLC fractions of dehydroxyxanthopterin B2 and co-purified compounds. **b**, Comparison of MS/MS fragmentation patterns for dehydroxyxanthopterin B2 and co-purified pteridines, with structures determined via HMBC and MS/MS. **c**, Structural overview of pteridines: a pyrazine ring (light blue) fused to a pyrimidine ring (dark blue). Pteridines are classified into lumazines (with a carbonyl group at C-2, red) and pterins (with an amino group at C-2, red). **d**, Summary of purified and identified lumazines (above) and pteridines (below). MF = molecular formula, EM = exact mass. **e**, Conversion of different pteridines to tetrahydrobiopterin, relevant enzymes are depicted in grey boxes: SPR = sepiapterin reductase, DHPR = dihydropteridine reductase, DHFR = dihydrofolate reductase.

### Purification and structure elucidation of the CutA binder dehydroxyxanthopterin B2

To determine the structure of the CutA ligand (compound 1, *m/z* 223.0824 [M+H]^+^), we scaled up the cultivation of *S. elongatus* WT and purified this compound via solid-phase extraction and HPLC. Growth of the culture and abundance of the putative CutA ligand are presented in **Supplementary Fig. 5**. In addition to compound 1, three nitrogen-containing heterocyclic compounds with similar MS/MS fragmentation patterns were also purified (**Fig. 3b**). Compounds 1 (*m/z* 223.0824 [M+H]^+^) and 3 (*m/z* 221.0675 [M+H]^+^) were detected at lower levels, while compounds 2 (*m/z* 222.0991 [M+H]^+^) and 4 (*m/z* 220.0831 [M+H]^+^) were produced at higher titers. Based on their similar masses and predicted molecular formulas, we hypothesized that elucidating the structure of any of these metabolites would reveal the architecture of this metabolite family.

Targeting compound 4, we were able to purify enough material for NMR analysis (**Supplementary Fig. 7-11**). The ^1^H NMR and HSQC NMR spectra of compound 4 in DMSO-d6 showed one aromatic proton (δ_H_ 9.08), two aliphatic protons (δ_H_ 3.13 (2H)), and one methyl group (δ_H_ 1.10 (3H)). The presence of three exchangeable protons was confirmed with HPLC-MS analysis using MeCN and D_2_O as eluents, wherein compound 4 became [M+D]^+^ at *m/z* 224.1078 (Δ -0.9 ppm). The HSQC data further revealed the carbon chemical shifts of the protonated carbons (δ_H_ 9.08/δ_C_ 147.5; δ_H_ 3.13/δ_C_ 29.8; δ_H_ 1.10/δ_C_ 7.5). A COSY correlation connected H3-3’ (δ_H_ 1.10) and H2-2’ (δ_H_ 3.13); shared HMBC correlations from H2-2’ and H3-3’ to C-1’ (δ_C_ 200.1) then outlined a propionyl moiety. Based on the propionyl substructure, molecular formula, and natural product database searches, 6-propionylpterin (exact mass: 219.0756; molecular formula: C_9_H_9_N_5_O_2_) was proposed as the structure of compound 4. The UV/vis data of compound 4 (λ_max_ 216, 304, 346 nm) was also consistent with that reported for 6-propionylpterin^37^. This pterin, found in different bacteria, is described as blue-fluorescent^38^ and is supposedly a cofactor for cyanide monooxygenase and involved in the degradation of cyanide^39^.

^1^H NMR data pertaining of compound 2 could be deciphered from ^1^H NMR data of compound 4/compound 2 mixtures (compound 2 slowly oxidized to compound 4), showing two heteroatom-substituted protons (δ_H_ 4.09), two aliphatic protons (δ_H_ 2.81 (^2^H)), and one methyl group (δ_H_ 0.98 (3H)). Four exchangeable protons could be observed (**Supplementary Fig. 11**). Again, using the putative propionyl substructure, molecular formula, and natural product databases, isosepiapterin was proposed as the structure of compound 2 (exact mass 221.0913, molecular formula: C_9_H_11_N_5_O_2_). The UV/vis data of compound 2 (λ_max_ 214, 264, 286 (sh), 410 nm) was also consistent with that reported for isosepiapterin, corroborating previous reports in *S. elongatus*, where absorption peaks at 270 nm and 410 nm were also observed for isosepiapterin^40^.

At this stage, the structures of compound 1 and compound 3 were identified as lumazines, corresponding to the pterin compounds 2 and compound 4, respectively. Specifically, compound 1 was determined to be a previously undescribed lumazine, which we named dehydroxyxanthopterin B2 due to its structural similarity to xanthopterin B2 (PubChem CID 439706, KEGG ID C02333), a metabolite previously identified in silkworms (*Bombyx mori*)^41^ and mouse cerebellum extracts^42^. Dehydroxyxanthopterin B2 has an exact mass of 222.0753 and a molecular formula of C_9_H_10_N_4_O_3_. The UV/Vis data of dehydroxyxanthopterin B2 (λ_max_ 216, 282, 398 nm) is consistent with its structure. Compound 3 was identified as 6-propionyllumazine, with an exact mass of 220.0596 and a molecular formula of C_9_H_8_N_4_O_3_. Its UV/Vis data (λ_max_ 214, 272, 326 nm) agreed with a previous study, which reported the presence of 6-propionyllumazine in the extract of a bioluminescent polychaete^43^ without describing its function. All identified compounds belong to the pteridine class of metabolites which are characterized by a heterocyclic pteridine ring structure (C6H_4_N_4_), consisting of a fused pyrimidine and pyrazine ring (**Fig. 3c**). These metabolites are further divided into pterins and lumazines. Pterins are distinguished by an amino group at the C-2 position, while lumazines possess a carbonyl group at C-2 (**Fig. 3c**). Pterins are well-documented as pigments, cofactors and redox mediators^44^ across biological systems, while lumazines remain understudied.

To confirm the interaction of pteridines with CutA, we performed a series of additional native MS measurements using a pre-purified HPLC fraction enriched in dehydroxyxanthopterin B2 (compound 1) along with co-purified compounds of similar mass, retention time, and polarity. This fraction was expected to have a higher overall pteridine concentration compared to the crude cell extract that was previously analyzed. With this data, and the obtained information about the spectral relationship of the compounds in the HPLC fraction, a network was generated showing the predicted compound classes (**Fig. 3a**). The network predominantly contained unannotated pteridines and their derivatives. We observed binding between CutA and masses corresponding to isosepiapterin, 6-propionylpterin, and dehydroxyxanthopterin B2.

Pterins exhibit multi-electron redox reactivity, mirroring the redox capabilities of transition metals^45^. They exist in multiple oxidation states—oxidized, semi-reduced (dihydro), and fully reduced (tetrahydro)—interconverting via two-electron, two-proton redox reactions. Many pterins, such as biopterin, show biological activity predominantly in their reduced tetrahydro form, such as tetrahydrobiopterin (BH_4_), a crucial cofactor in various biological processes^46,47^. **Figure 3e** illustrates the stepwise reduction of sepiapterin to dihydrobiopterin, followed by its further conversion to tetrahydrobiopterin (BH_4_) through two-electron, two-proton redox reactions. Based on this knowledge, we hypothesized that our purified compounds 1-4 may be redox equivalents of each other (**Fig. 3d**). Due to its central role in cellular processes, we focused on tetrahydrobiopterin and its oxidized form, 7,8-dihydrobiopterin (BH_2_) to elucidate the potential functions of CutA. To investigate this, we obtained both compounds and tested their binding to CutA.

### Verification of CutA interactions with pteridines and copper binding

To investigate the interaction of *E. coli* CutA with purified pteridine derivatives, we conducted further native mass spectrometry, microscale thermophoresis (MST), and nano differential scanning fluorimetry (NanoDSF).

Native mass spectrometry experiments revealed that CutA forms complexes with purified pteridines, observed by mass shifts between apo and ligand-bound peaks (**Fig. 4a, left**). For some pteridines, especially for dehydroxyxanthopterin B2 (DXan B2), we saw more than one shift, indicating multiple binding events (**Supplementary Fig. 7**). NanoDSF assays demonstrated that BH_4_ and BH_2_ significantly destabilize CutA, decreasing the melting temperature (Tm) of the protein by 11.5 °C and 8.2 °C, respectively. A decrease in Tm was also observed for DXan B2; however, it could not be quantified due to limitations in the concentration range available for testing (**Fig. 4a, right**). Titration experiments further confirmed concentration-dependent binding of BH_2_, isosepiapterin, and DXan B2, while folic acid, which contains a pteridine moiety, showed no significant interaction (**Fig. 4b**).

**Figure 4:**
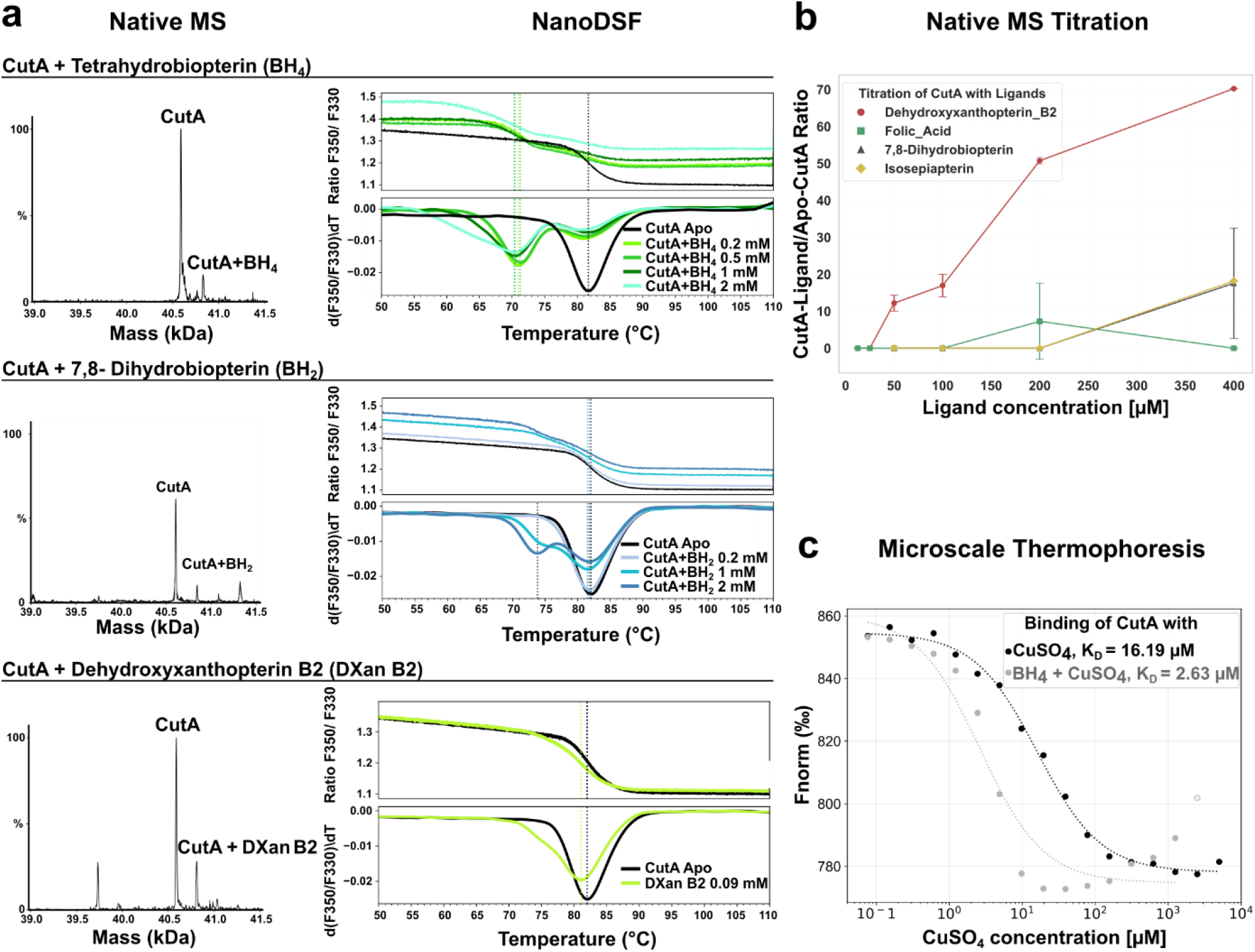
Orthogonal Binding Studies of Pteridines and CuSO_4_ to CutA Using Native MS, NanoDSF, and MST. **a**, Binding of BH_4_, BH_2_ and DXan B2 Left: Native MS spectra show mass shifts corresponding to the formation of CutA-pteridin complexes with BH_4_, BH_2_, and DXan B2. The corresponding NanoDSF measurements are depicted on the right, showing changes in CutA thermal stability (Tm) upon ligand binding. **b**, Native MS titration curves indicate concentration-dependent interactions of CutA with various pteridines (duplicates; isosepiapterin: single measurement) **c**, MST measurements reveal binding affinities of CuSO_4_ to CutA. Black curve: without BH_4_; grey: with constant 0.5 mM BH_4_ with dissociation constants (Kd) of 16.19 µM or 2.63 µM, respectively.

Given conflicting reports about the ability of *E. coli* CutA to bind copper ions^3,4,48,49^, we aimed to further explore this interaction. Viability assays in liquid cultures without CuSO_4_ showed no difference between WT and mutant under standard growth conditions (see **Supplementary Fig. 8**,**9**). We performed MST experiments to confirm binding of copper ions to CutA and also to evaluate whether pteridines can influence the binding of copper sulfate (CuSO_4_). Binding curves (**Fig. 4c**) demonstrated CuSO_4_ binding to CutA with a K_D_ of about 16.19 µM. In a second experiment, CutA was incubated with 0.5 mM BH_4_, and CuSO_4_ binding was measured by MST. The data suggested a reduced K_D_ value of 2.63 µM in the presence of BH_4_, indicating a potential role for pterin as a cofactor of CutA in copper binding. Many enzymes rely on both a metal ion and a pterin derivative as cofactors for essential biochemical reactions, including redox processes. Molybdopterin-dependent enzymes like nitrate reductase (EC1.7.1.2), xanthine oxidase (EC1.17.3.2)^50,51^, and sulfite oxidase (EC1.8.3.1)^52^ require molybdenum, while tetrahydrobiopterin-dependent hydroxylases, such as phenylalanine hydroxylase (EC1.14.16.1)^53^, tyrosine 3-monooxygenase^54^ (EC1.14.16.2) or tryptophan hydroxylase (EC1.14.16.4) utilize iron. Tetrahydrobiopterin has been shown to reduce the Mn(III) center of a porphyrin to Mn(II)^55^ and is also known to form complexes with cupric ions, facilitating the reduction of Cu(II)^45^ (see also **Supplementary Fig. 8**). This dual reliance on metal ions and pterin derivatives raises intriguing questions about the potential role of CutA in mediating such interactions. Additionally, pteridine biosynthesis appears to be influenced by metal availability. The *pts* gene, which encodes 6-pyruvoyl tetrahydrobiopterin synthase—a zinc-dependent enzyme^56^ that structurally resembles CutA in its structure and is involved in the early biosynthesis of pteridines—was upregulated in cyanobacteria under both iron and copper perturbations^56^. Given the high levels of pteridine glycosides produced by cyanobacteria and the role of pterins in stress responses to UV and blue light^57^, it is plausible that CutA’s interaction with pterins extends beyond copper binding, potentially contributing to broader metal stress adaptation mechanisms. While further experiments are required to confirm and assess the mechanisms of pterin-mediated copper binding, our results provide new functional insights into CutA and the role of pterins in cellular copper homeostasis.

## Discussion

Our study uncovered the interaction between the conserved trimeric protein CutA and pteridine metabolites, including tetrahydrobiopterin. Using a combination of native MS and NanoDSF, we show that these interactions destabilize CutA, suggesting a functional link between this protein and pterin redox chemistry. Given the critical roles of pterins in redox regulation, enzymatic catalysis, and oxidative stress responses, these findings provide new insights into CutA’s physiological role. While CutA has been primarily associated with copper tolerance, our data points to a more nuanced role for this protein, potentially bridging copper homeostasis and cellular redox regulation via pteridine metabolism. This function could be particularly relevant in mitigating oxidative stress caused by copper dysregulation, a process that threatens critical pathways such as respiration and photosynthesis. *Dehydroxyxanthopterin B2*, a novel pteridine with strong binding affinity for CutA, further underscores the evolutionary conservation and functional versatility of pteridine metabolites across biological systems. Our results suggest that CutA plays a key role in cellular stress responses, particularly in mitigating reactive oxygen species generated by heat and ampicillin-induced stress. Under these conditions, we observed metabolic imbalances, suggesting that CutA may help to maintain cellular integrity by regulating copper and redox homeostasis. While further research is required to uncover the broader implications of CutA-pteridine interactions, our study advances the understanding of CutA beyond its structural resemblance to PII proteins.

In summary, our data identified pteridines as CutA binding metabolites, which increase its copper affinity. Together these results render CutA as a potential mediator of redox biology and stress response mechanisms, particularly in the context of copper-associated oxidative stress.

## STAR Methods

### Growth conditions

*Synechococcus elongatus* precultures were cultivated in shaking flasks containing BG11 medium according to Rippka et al.^58^ with additional 5 mM NaHCO_3_ under photoautotrophic conditions with continuous light (30-60 µmol photons m^-2^s^-1^E) at 28 °C and constant shaking (120 rpm), if not described differently. *E. coli* precultures were cultivated in LB broth at 37 °C with constant shaking (250 rpm).

### Drop Dilution Assay

*S. elongatus* culture (OD_750_ = 1) was serially diluted (10°–10^−6^) in BG11 medium. Subsequently, 5 µL of the dilutions were dropped in biological replicates on BG11 agar plates, which were incubated at 28 °C with continuous illumination (30-60 µmol photons m^-2^ s^-1^E).

### Ampicillin treatment of *S. elongatus*

50 mL early exponential culture (OD_750_ of 0.4 in BG11) were treated with 10 µg/mL ampicillin and incubated with shaking at 28 °C with constant illumination (30-60 µmol photons m^-2^ s^-1^E) for two weeks.

### Bacterial strains and molecular cloning of seamless *E. coli ΔcutA* mutant

*Synechococcus elongatus* PCC 7942 wildtype (WT) and *S. elongatus ΔcutA::kan* strain, as reported in Selim (2021)^4^, were used in this study. *E. coli* strains included the *E. coli* BW25113 wild-type (WT) and a *ΔcutA* mutant was cloned for this project. The *cutA* deletion mutant without an antibiotic resistance cassette was created using scarless genome editing based on the method described by Kim et al.^59^. The deletion mutant was verified through PCR and sequencing to confirm the deletion of the *cutA* gene. Fragments were sequenced through the “GATC LIGHTRUN” service by Eurofins Scientific (Luxembourg), and the results were analyzed using Clone Manager 9 (Scientific & Educational Software, Denver, CO, USA). Sequence alignment is shown in **Supplementary Fig. 18**.

### Protein expression and purification

Proteins were expressed using the pASK-IBA vector (IBA Lifesciences) and purified on an AEKTA FPLC purifier system using StrepTrap HP 5 mL columns (Cytiva), following the respective manufacturers’ protocols. For the induction of CutA expression, *E. coli* Lemo^60^ strains harboring pASK [*E. coli*_*cutA1*] or pASK[*S. elongatus*_*cutA*] were grown in double-concentrated lysogeny broth (LB) supplemented with ampicillin (100 µg/mL) and chloramphenicol (34 µg/mL). Cultures were incubated at 37 °C with shaking (120 rpm) until reaching an OD_600_ of 0.5. CutA expression was induced by adding anhydrotetracycline to a final concentration of 200 µg/L, followed by incubation at 20 °C with shaking (120 rpm) for 12–14 hours. Cells were harvested by centrifugation (4000 × g, 12 min, 4 °C), and the resulting pellets were resuspended in lysis buffer (100 mM Tris-HCl, pH 8.0, 150 mM NaCl, 1 mM EDTA) containing DNase I, RNase A, lysozyme, and protease inhibitors. Cell disruption was achieved by sonication, and debris was removed by centrifugation (39,000 × g, 60 min, 4 °C) followed by sterile filtration (0.22 µm filters). Proteins were eluted with an elution buffer (100 mM Tris-HCl, pH 8.0, 150 mM NaCl, 1 mM EDTA, 5 mM d-desthiobiotin). Protein quality was assessed by SDS-PAGE, and collected fractions were concentrated using Amicon Ultra Centrifugal Filters (3 kDa cutoff, Merck-Millipore). The buffer was exchanged to a dialysis buffer (20 mM Tris-HCl, pH 7.8, 150 mM KCl, 50% glycerol), and the concentration of CutA was determined using bicinchoninic acid (BCA) or Bradford assay. SDS pages and chromatogram of purification are shown in **Supplementary Fig. 17 and 18**.

### Targeted metabolomics

Cells were cultivated in minimal medium (*S. elongatus* PCC 7942 in BG11, *E. coli* in M9 medium) to an OD_750_ of ∼0.45. A sample volume of 2 mL (*E. coli*) or 10 mL (*S. elongatus*) was collected and filtered through 1.2 µM pore-size filters (WHA1822025, Cytiva, Marlborough, MA, USA). Filters were transferred into 500 µL of acetonitrile: methanol: H_2_O (40:40:20) at −20 °C and incubated overnight in the extraction solvent at the same temperature. After cell lysis, the extract was transferred to a new tube and centrifuged. To ensure proper lysis of cyanobacterial cells, the *S. elongatus* extract was transferred into tubes containing glass beads and subjected to ribolysis at 6.5 m/s for 30 seconds in 2 cycles, with a 5-minute break between cycles, then centrifuged at >13,000 × g for 15 minutes at −9 °C. The supernatant was transferred to a new tube and stored at −80 °C until analysis. LC-MS/MS analysis was performed using a 6495A triple quadrupole LC/MS system (Agilent, Santa Clara, CA, USA), with relative quantification achieved using a _13_C internal standard. Detailed measurement and analysis parameters are described in Guder et al._61_. For data analysis, the ratios of the _12_C sample to the _13_C internal standard were calculated and normalized to cell density. Next, fold-change of each sample relative to the mean of the controls (WT) was determined and log_2_-transformed for improved visualization (log_2_ FC = 0: no change; log_2_ FC = 1: 100% increase; log_2_ FC = -1: 50% decrease).

### Non-targeted metabolomics

Five WT and five *ΔcutA::kan* strains were cultivated in pentaplicates in 200 mL BG11 medium in 500 mL shaking flasks under continuous light (∼50 µE) at 28 °C. Half of the cultures were harvested during exponential growth (OD_750_ ∼0.6) and the other half after 14 days in stationary phase. Cultures were centrifuged (4200 rpm, 60 min, 4 °C), and 75 mL per culture were harvested. Cell pellets were frozen in liquid nitrogen and stored at - 20 °C. The growth medium (BG11) was treated similarly as a control. Metabolites were extracted by adding 10 mL of 20% MeOH per gram of wet cell mass, vortexed, and subjected to ultrasonic bath treatment for 10 minutes. Cell extracts were dried, normalized to 2.5 mg/mL, and prepared for LC-MS measurement. LC-MS/MS data were collected using a Vanquish ultrahigh-performance liquid chromatography (UHPLC) system coupled to a Q Exactive HF mass spectrometer (Thermo Fisher Scientific, Bremen, Germany) with heated electrospray ionization (HESI). Chromatographic separation was performed at a constant flow rate of 0.5 mL/min with a mobile phase consisting of A (H_2_O + 0.1% formic acid) and B (MeCN + 0.1% formic acid), using a gradient from 5% to 99% mobile phase B. 0.0 min: 5% B, 8.0 min: 50% B, 10.0 min: 99% B, 13.0 min: 99% B, 13.1 min: 5% B. The separation was achieved using a Kinetex 1.7 µm C18 reversed-phase UHPLC column (100 Å pore size, 50 × 2.1 mm l x i.d.) by Phenomenex (Torrance, California, USA). Data were acquired in data-dependent acquisition (DDA) mode, selecting the five most abundant ions for fragmentation following each MS survey scan (Top5).

### Native MS and micro-flow LC-MS/MS data acquisition

Native mass spectrometry (MS) of CutA with ligands or cell extracts was performed using a Q Exactive HF Orbitrap instrument (Thermo Fisher Scientific). Purified proteins (2 mg/mL) were rebuffered into 10 mM ammonium acetate (pH 8) through five cycles of ultrafiltration (12,000 × *g*, 12 min, 4 °C) using centrifugal filters. Data acquisition for native MS and micro-flow LC-MS/MS followed the protocol by Reher et al., 2022^24^. To stabilize pH, 100 mM ammonium acetate buffer (150 µL/min) was delivered post-column via a make-up pump. Proteins were injected at 2 µL/min using an integrated syringe pump, while 5 µL of cell extracts were introduced at 100 µL/min. The MS scan range was set to 2500–4000 *m/z* at a resolution of 240,000 (*m/z* 200). Chromatographic separation was performed on a Phenomenex Kinetex EVO C18 column (1.7 µm, 100Å, 150 × 1.0 mm l. x i.d.). All-Ion Fragmentation (AIF) was performed with a collision energy (CE) of 10 eV to fragment ions across the full *m/z* range. Native MS measurements of cell extract of *S. elongatus* and *E. coli* (WT and *ΔcutA* strains) were carried out after the cells were extracted with 20% or 80% MeOH to obtain a broad range of polar or nonpolar compounds (**Supplementary Fig. 8**). The cell extracts were dissolved in 50% MeOH and separated using reversed-phase Ultra High-Performance Liquid Chromatography (UHPLC). Metabolomics runs were conducted under identical chromatographic conditions, excluding protein infusion. Sample volumes of 5 µL were injected at a flow rate of 100 µL/min, using data-dependent acquisition (DDA, Top 5) with a scan range of 80– 1500 *m/z*. Pteridine titrations were performed using a 5–99% gradient over 5 minutes. Native mass spectrometry (MS) was carried out in the 2000–5000 *m/z* range, with a resolution of 240,000 at *m/z* 200 and positive polarity and collision energy of 10 eV. A 33% ammonium acetate solution was infused via a make-up pump. Ligands dissolved in 50% MeOH were injected as 2 µL aliquots. For the binding curve, pteridines were prepared at 800 µM stock concentrations and diluted 1:2 for binding assays. Native MS spectra were deconvoluted using UniDec^62^ (Charge range 6-20, mass range 10000-50000, mass sampling every 1.0 Da, peak detection range 5.0 ppm, peak detection threshold 0.1), and shifted mass peak intensities were expressed as percentages relative to the apo protein.

### LC-MS/MS data analytics workflow

Feature tables (.csv) and MS/MS spectra files (.mgf) were deposited in the MassIVE repository. LC-MS/MS raw files were converted to centroided mzML format using MSConvert from the ProteoWizard software package^63^. The files were then processed using MZmine3^64^ for feature detection and alignment. The LC-MS/MS data were further submitted to GNPS^65^ and GNPS2^66^ (gnps2.org) for feature-based molecular networking analysis and metabolite annotation. The obtained metabolomics data were further processed and analyzed using SIRIUS^67^ to predict molecular formulas and annotate fragmentation patterns from the high-resolution MS data, which provided further insights about the metabolites in the HPLC fraction. Feature-based molecular networking results were screened and visualized in Cytoscape^68^, and statistical analyses were carried out using the Statistical Analysis of Feature-Based Molecular Networking script^69^ or the associated web application (https://fbmn-statsguide.gnps2.org). Background noise was eliminated by removing blanks with a cutoff of 0.3 to improve data accuracy. To examine relationships between samples, Principal Coordinates Analysis (PCoA) was performed using the Canberra distance matrix to visualize dissimilarities among the samples.

### Growth conditions for compound purification

*Synechococcus elongatus* PCC 7942 was grown under continuous light conditions in a 10 L flask containing BG11 medium and, in parallel, for the purpose of higher cell mass in a CellDEG® (CellDEG GmbH, Germany) system with freshwater medium.

### Pteridine purification

Cells were harvested by centrifugation (4200 rpm, 4 °C). The cell pellets were extracted with 20% MeOH. The cell extract was dry-loaded onto 5x Celite and fractionated using silica gel (100% hexane, 30% EtOAc in hexane, 100% EtOAc, 1% MeOH in DCM, 10% MeOH in DCM, 100% MeOH, each fraction ∼5 column volumes). The 10% and 100% MeOH fractions were combined and concentrated. The product was first purified via HPLC [Kinetex C18, 250 x 21.2 mm, 5 µm, 100 Å pore size, 5% MeCN in water (0.1% TFA) for 5 min, 5-30% MeCN (0.1% TFA) in water over 15 min, 13 mL/min, 254 nm, tR = 14-20 min]. The product was further purified via two rounds of isocratic HPLC [Luna C18(2), 250 x 10 mm, 5 µm, 100 Å pore size, 10% MeCN in water (0.1% TFA), 3 mL/min, 254 nm, tR = 14-16 min; then PFP C18(2)column 250 x 10 mm, 5 µm, 100 Å pore size, 10% MeCN in water (0.1% TFA), 3 mL/min, 254 nm, tR = 16-18 min] to yield <1 mg of compound 4 and <1 mg of a mixture of compound 2 and 4.

### Structure elucidation of compounds 1-4

^1^H and 2D (COSY, HSQC, HMBC, NOESY) NMR spectra were recorded on a Bruker Avance III HDX 700 MHz spectrometer fitted with a 5 mm Prodigy (^1^H,^19^F/^13^C/^15^N) TCI Cryoprobe. ^1^H NMR data were recorded at 700 MHz in DMSO-d_6_ (2.50 ppm), and ^13^C NMR data were recorded at 175 MHz in DMSO-d_6_ (39.5 ppm). NMR spectra were processed using MestReNova (Mnova 14.3.0, Mestrelab Research) software. The following abbreviations are used to indicate the multiplicity in ^1^H NMR spectra: s = singlet, d = doublet, t = triplet, q = quartet.

1. λ_max_ (MeCN/H_2_O/0.1% FA): 216, 282, 398 nm; HRESIMS *m/z* [M+H]^+^ 223.0831 calcd for C_9_H_10_N_4_O_3_, Dehydroxyxanthopterin B2.
2. λ_max_ (MeCN/H_2_O/0.1% FA): 214, 264, 286 (sh), 410 nm; HRESIMS *m/z* [M+H]^+^ 222.09910, calcd for C_9_H_11_N_5_O_2_, Isosepiapterin ^1^H NMR: δ 4.09 (s, 2H), 2.81 (q, J = 7.5 Hz, 2H), 0.98 (t, J = 7.5 Hz, 3H)
3. λ_max_ (MeCN/H_2_O/0.1% FA): 214, 272, 326 nm; HRESIMS *m/z* [M+H]^+^ 221.0674 calcd for C_9_H_8_N_4_O_3_, 6-Propionyllumazine
4. λ_max_ (MeCN/H_2_O/0.1% FA): 216, 304, 346 nm; HRESIMS *m/z* [M+H]^+^ 220.08345, calcd for C_9_H_9_N_5_O_2_, 6-Propionylpterin

^1^H NMR: δ 9.08 (s, ^1^H), 3.13 (q, J = 7.3 Hz, 2H), 1.10 (t, J = 7.3 Hz, 3H); HSQC/HMBC: δ_H_ 9.08/δ_C_ 147.5; δ_H_ 3.13/δC 29.8; δ_H_ 1.10/δ_C_ 7.5; δ_C_ 200.1

### NanoDSF (differential scanning fluorimetry)

Nanoscale differential scanning fluorimetry (nanoDSF) was performed using a Prometheus NT.48 instrument (Nanotemper Technologies, Munich, Germany) with standard Prometheus capillaries (PR-C002). Protein samples and ligands, both dissolved in HEPES buffer, were analyzed with a temperature ramp of 0.5 °C/min from 30 °C to 110 °C. Thermal stability was assessed by monitoring the fluorescence ratio (F_350_/F_330_) of tryptophan residues, with the melting temperature (Tm) determined from the inflection point of the fluorescence ratio curve and confirmed by first derivative analysis. ∼300 µM of *E. coli* CutA trimer was used for the measurements.

### MST (Microscale Thermophoresis)

To investigate protein interactions via MicroScale Thermophoresis (MST), *E. coli* CutA was labeled with a fluorescent dye using the Protein Labeling Kit RED-NHS® 2nd Generation (NanoTemper Technologies). NHS labeling attaches fluorescent dyes to proteins by reacting with primary amines to form stable amide bonds. This was necessary due to the fluorescence properties of pteridines. MST measurements were conducted with 10 nM protein concentration.

The protein and ligands were dissolved in HEPES or MST buffer (50 mM Tris, 150 mM NaCl, 0.05% Tween-20, 10 mM MgCl_2_, pH 7.4) for analysis. Ligands were prepared in a titration series with 1:2 dilutions, starting at 2.5 or 5 mM, and loaded into Monolith NT.115 capillaries. MST was performed using the Monolith NT.115 with settings of 25 °C TempControl, 80% LED Power, and 20% MST Power. Dose-response data were analyzed with MO.Affinity Analysis Software v2.3 (NanoTemper) and further fitted using a custom Python script applying non-linear regression with a one_site_binding model.

## Data availability

Native MS with crude cell extracts: (MassIVE MSV000088919), Zenodo: https://doi.org/10.5281/zenodo.10171306 Native MS with HPLC extract of pteridine purification: MassIVE MSV000092597 Native MS of S. elongatus CutA + Cell extract or HPLC extract: MassIVE MSV000092597 Non-targeted Metabolomics of *S. elongatus* WT vs. *ΔcutA::kan* exponential and stationary growth phase (for PCoA): MassIVE MSV000092427 or Zenodo DOi: https://doi.org/10.5281/zenodo.10041900 Native MS CutA and different pteridines: Zenodo: https://doi.org/10.5281/zenodo.14749948, Zenodo. https://doi.org/10.5281/zenodo.14750590

## Acknowledgements

We thank Paolo Stincone, Michael Haffner, Nelli Deobald and Andreas Kulik for technical support as well as Prof. Dr. apl. Klaus Hantke and Gulliver Black for insightful discussions. We acknowledge funding through the Cluster of Excellence (EXC2124–390838134, Controlling Microbes to Fight Infections, CMFI) at the University Tuebingen (projects: 1-06.009 and 1-06.010). We are grateful to Dr. Libera Lo Presti for their critical review of the article.

## Author contributions

BCW, KF, and KAS conceptualized and planned physiology experiments. BCW, EN and AS performed cloning and phenotyping experiments. BCW and HPG performed heterologous expression and protein purification. BCW and DP conceptualized and planned the LC-MS/MS and native MS experiments. BCW, CG and DP conducted untargeted and native metabolomics experiments. JR, BCW and HL designed, conducted, and analyzed targeted metabolomics experiments. BCW and CCH purified pteridines. CCH analyzed NMR spectra and elucidated the structures. BCW and RA conducted NanoDSF and MST experiments under MDH’s supervision. BCW, CCH, and DP wrote the manuscript. All authors edited and approved the final version.

