## Supplementary_information for "Native metabolomics identifies pteridines as CutA ligands and modulators of copper binding"

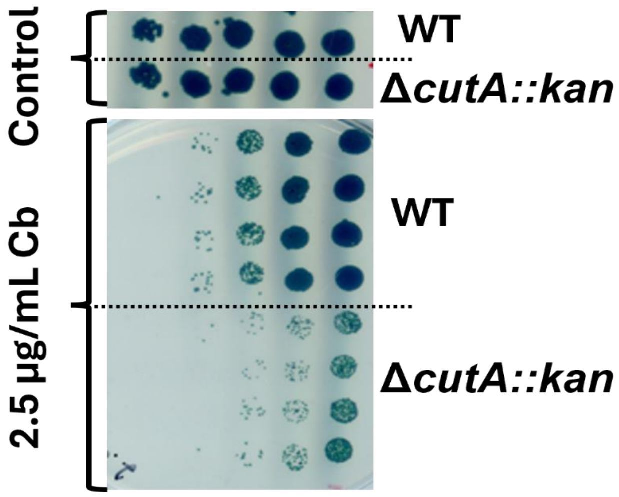

**Supplementary Figure 1: Viability assay of *Synechococcus elongatus* WT and *ΔcutA::kan* on BG11 agar supplemented with 2.5 µg/mL carbenicillin (Cb).** A drop dilution assay was performed with 1:10 serial dilutions, starting at an $OD_{750}$ of 1. Plates were incubated at 28°C under continuous light (30–60 µmol photons $m^{-2}s^{-}E^{1}$) for 14 days.

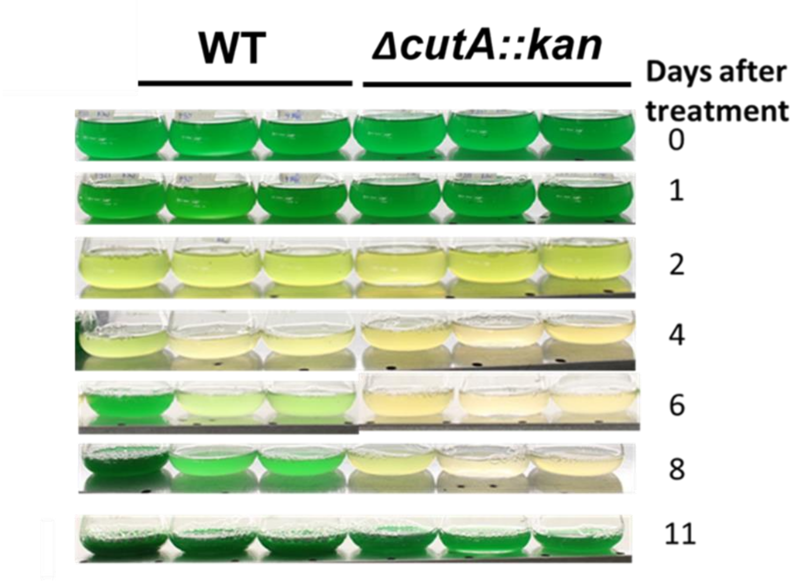

**Supplementary Figure 2: Repetition of viability assay in liquid cultures of *Synechococcus elongatus*** Cultures were treated with 10 µg/mL ampicillin and inoculated at a starting $OD_{750}$ of 0.4. Incubation was carried out under continuous light (30-60 µmol photons $m^{-2}s^{-1}E$) at 28°C in biological triplicates.

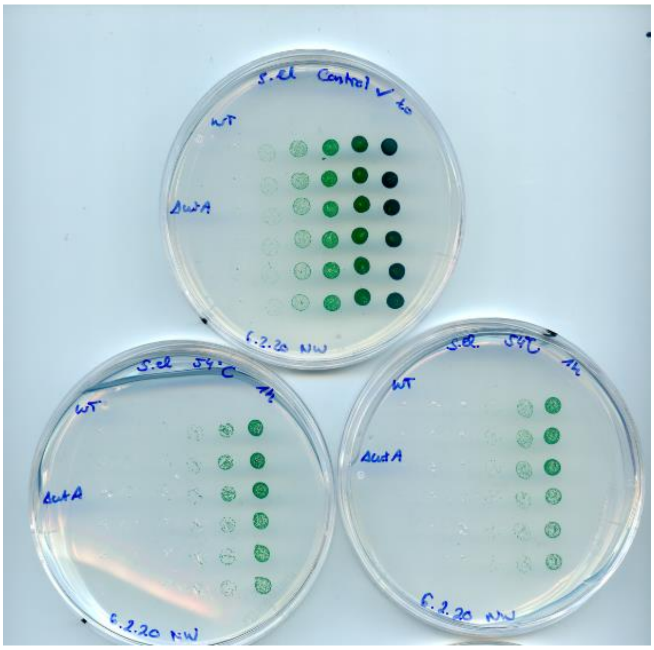

**Supplementary Figure 3: Full plates and repetition of the viability assay presented in Fig. 1.** The viability drop dilution assay demonstrates impaired recovery of the *ΔcutA::kan* mutant following heat stress. *Synechococcus elongatus* WT and *ΔcutA::kan* cultures were exposed to 50°C for 1.5 hours, serially diluted (1:10, right to left), and incubated at 28°C under constant light (30–60 µmol photons $m^{-2}s^{-}E^{1}$) for 7 days.

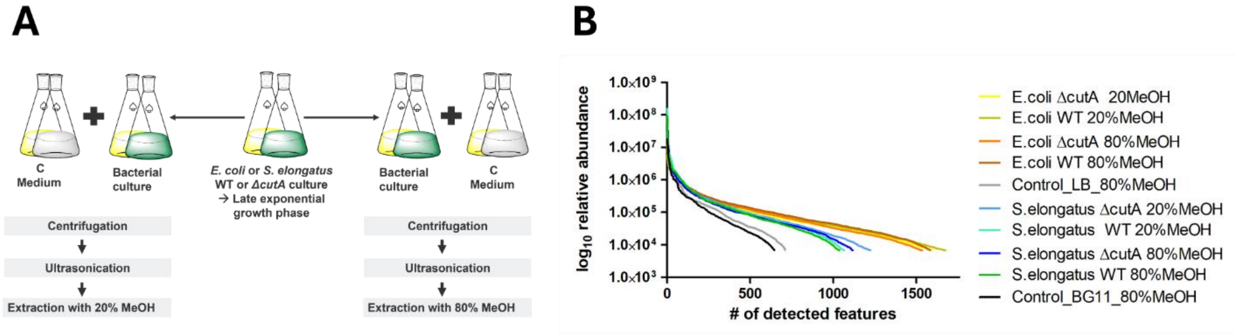

**Supplementary Figure 4: A) Extraction of *S. elongatus* and *E. coli* cells and B) metabolite abundance.** Extraction from late exponential phase cultures using 20% or 80% MeOH to capture a broad range of polar and non-polar compounds. B: Feature abundance in media (gray) compared to *E. coli* (yellow) and *S. elongatus* extracts (blue and green).

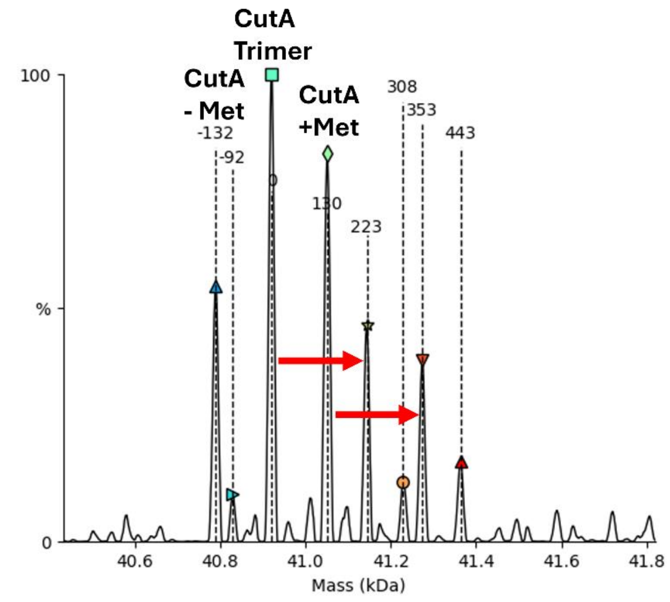

**Supplementary Figure 5: Native mass spectrometry analysis of *S. elongatus* CutA.** Testing of HPLC extract containing various pteridines, including dehydroxyxanthopterin B2. Native MS revealed three distinct CutA trimers, likely resulting from methionine aminopeptidase activity, as the peaks were spaced by the mass of one methionine. Data were deconvoluted using UniDec, revealing mass shifts corresponding to *Synechococcus elongatus* CutA in complex with dehydroxyxanthopterin B2 (shift indicated by red arrow).

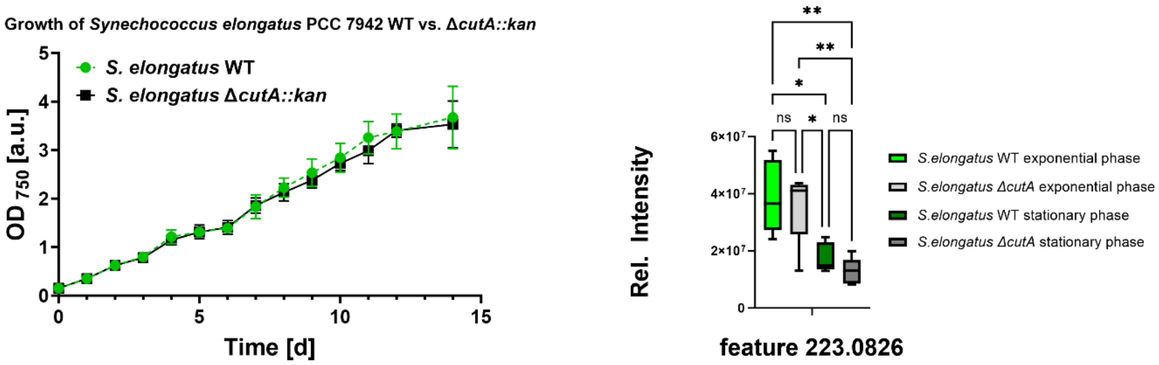

**Supplementary Figure 6: Growth and abundance of DXan-B2** Left: Growth of *S. elongatus* WT and *ΔcutA::kan* and right: abundance of dehydroxyxanthopterin B2. Statistical analysis was conducted using one-way ANOVA, followed by Tukey's multiple comparison test for post-hoc analysis. Analyses were performed in GraphPad Prism version 10.1.2 (334).

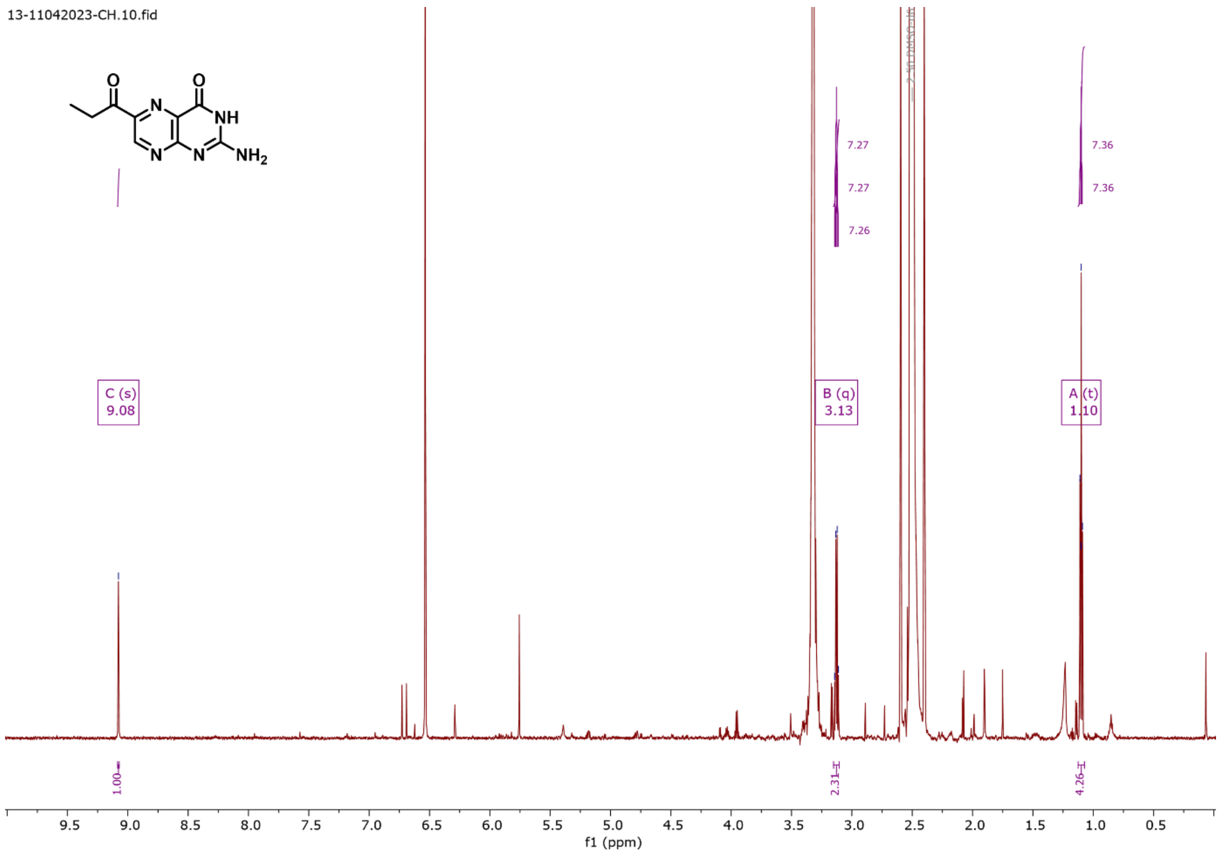

**Supplementary Figure 7: $^{1}$H NMR (DMSO-$d_6$, 700 MHz) of compound 4**

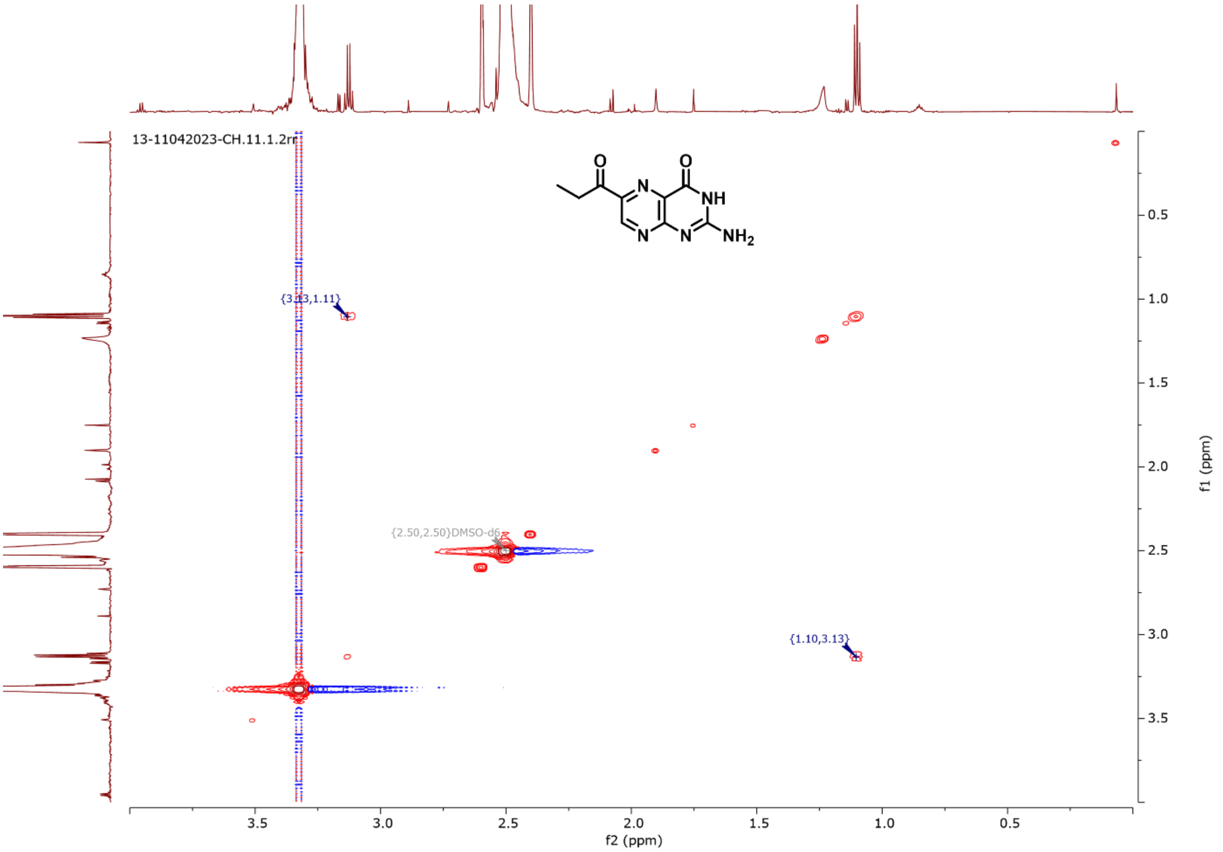

**Supplementary Figure 8: COSY (DMSO-d6, 700 MHz) of compound 4**

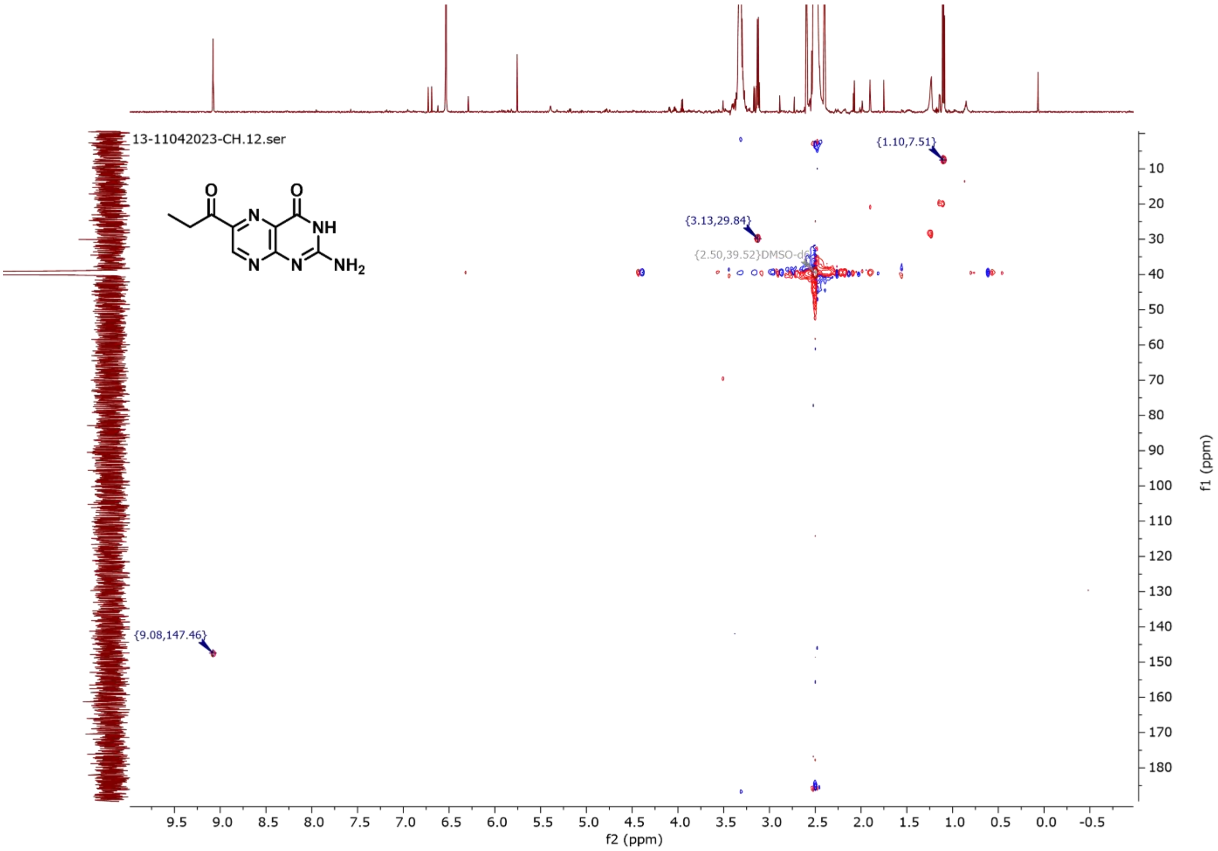

**Supplementary Figure 9: HSQC (DMSO-d6, 700 MHz) of compound 4**

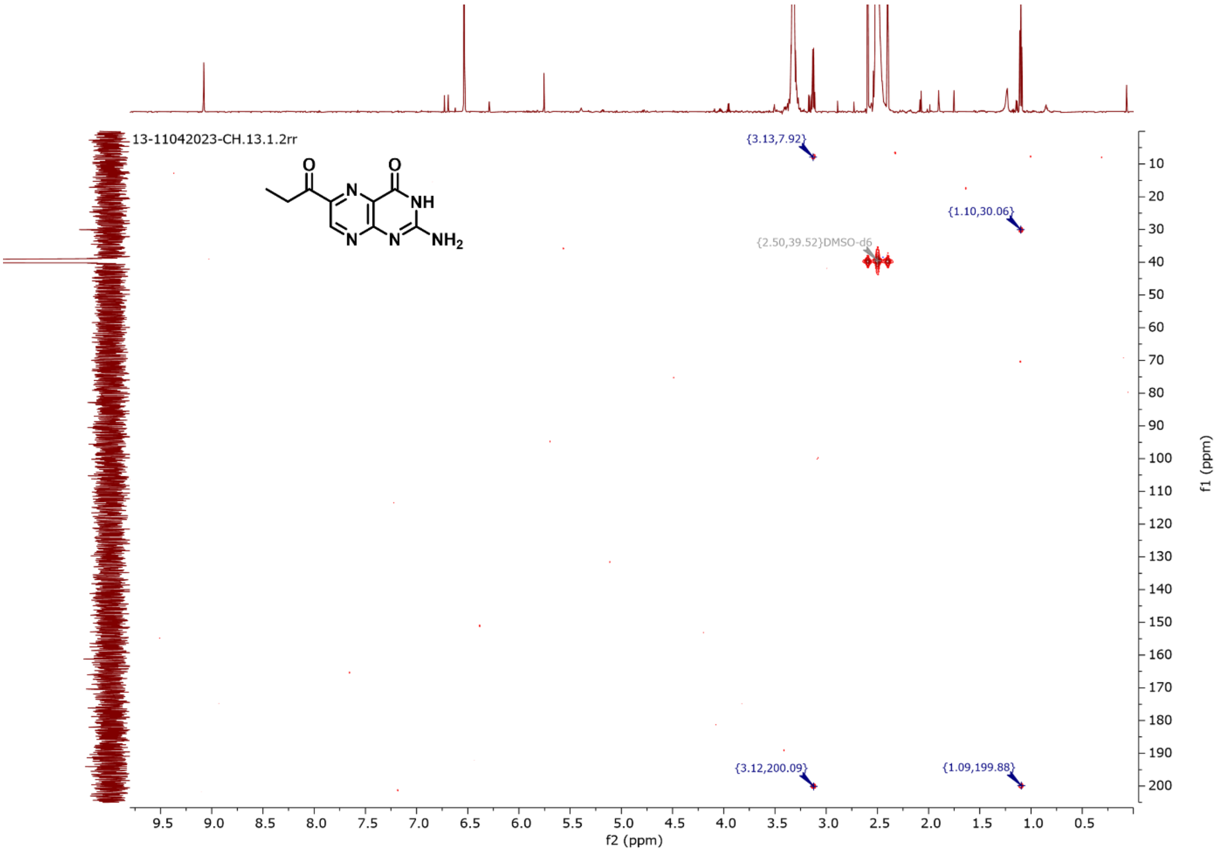

**Supplementary Figure 10:HMBC (DMSO-d6, 700 MHz) of compound 4**

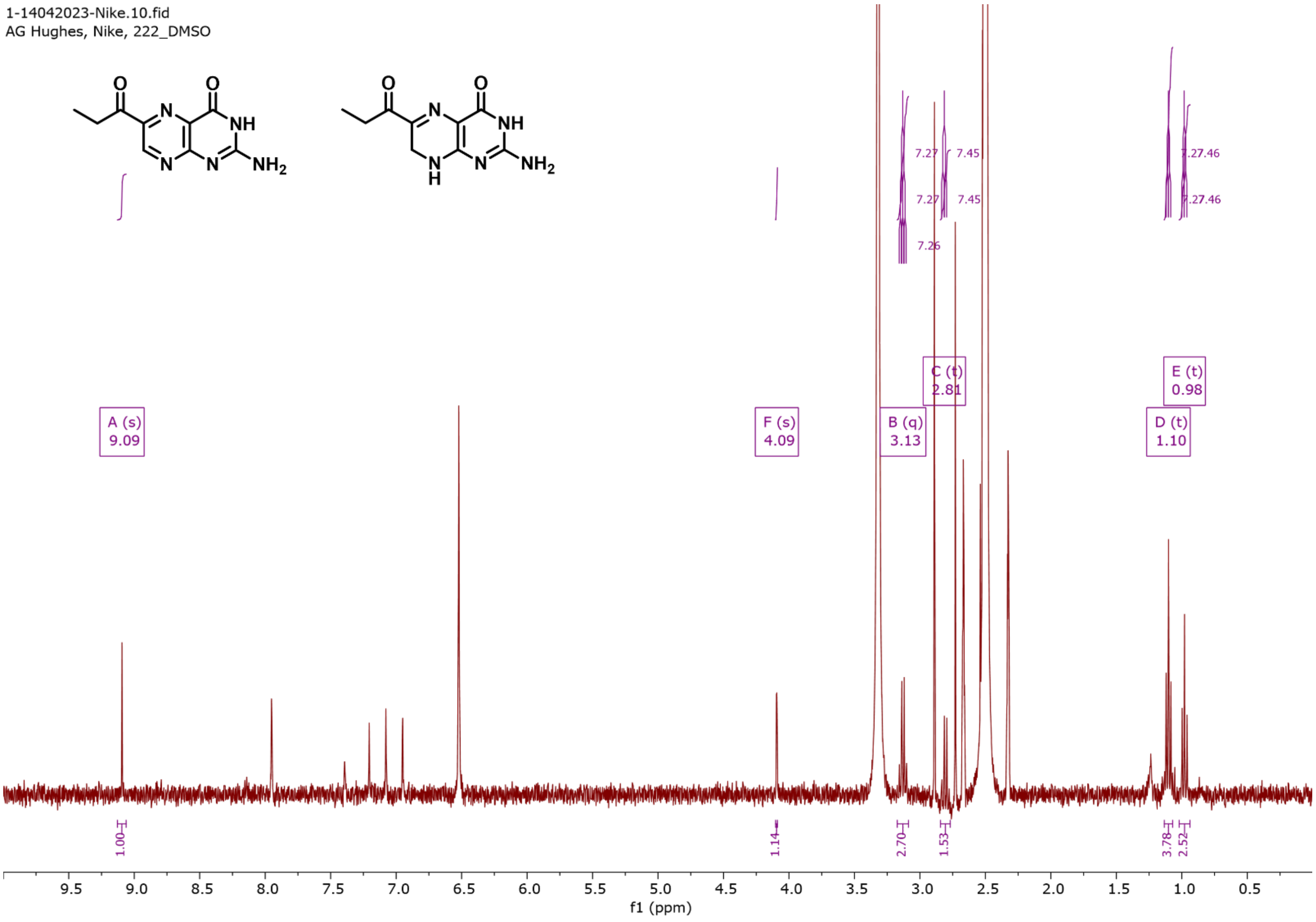

**Supplementary Figure 11:1H NMR (DMSO-d6, 700 MHz) of 2 and 4**

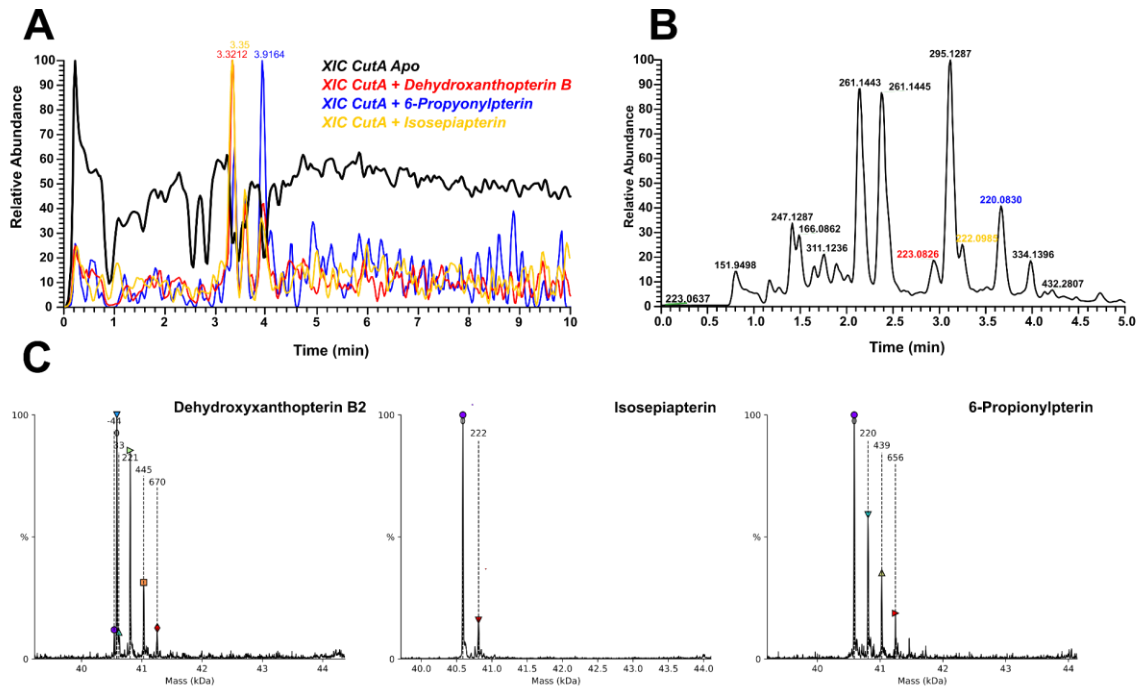

**Supplementary Figure 12: Different pteridines binding to CutA.** A) XIC of Apo E. coli CutA (black), XIC of CutA+ different identified pteridines in HPLC fraction B) TIC of HPLC fraction containing dehydroxyxanthopterin B C) Native MS of CutA with various pteridines, indicating potential multiple binding events.

#### Copper starvation of *S. elongatus* Wt and *ΔcutA::kan*

Given our finding that CutA binds copper, we hypothesized that CutA may function as a copper storage protein. To test this hypothesis, we initially grew cultures in BG11 medium, both with and without the addition of $CuSO_4$ (SI-Fig.13). Observing no significant differences in growth under these conditions, we introduced triethylenetetramine, a copper chelator that binds free copper ions with high affinity, to ensure the complete removal of any residual copper from the medium.

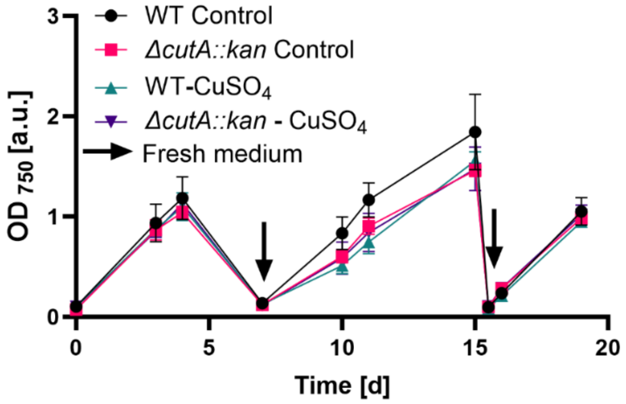

**Supplementary Figure 13: Effect of copper starvation on growth of *S. elongatus* WT and *ΔcutA::kan*** *Synechococcus elongatus WT and ΔcutA::kan strains were grown in copper-free BG11 medium (BG11 and BG11 without $CuSO_4$) in 50 mL plastic flasks. Cultures were inoculated at an initial $OD_{750}$ of 0.1, illuminated at 30-60 µmol photons $m^{-2}s^{-1}$E, and shaken at 120 rpm. All experiments were performed in biological triplicates.*

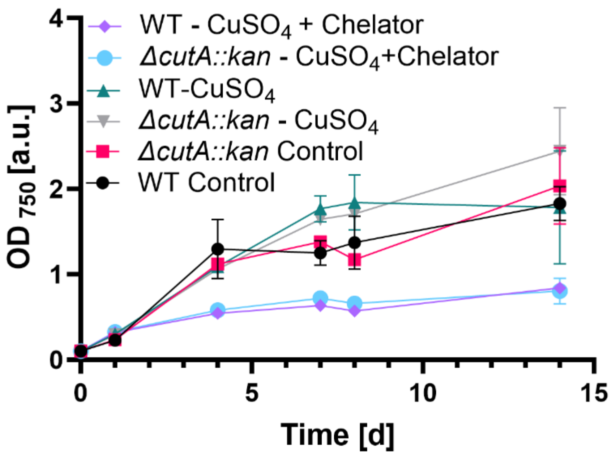

**Supplementary Figure 14: Effect of copper starvation with the chelating agent triethylenetetramine on growth** *Cultures were inoculated at an initial $OD_{750}$ of 0.1 and grown under the following conditions: 2 replicates with copper (+Cu), 3 replicates without copper (-Cu), and 2 replicates without copper but with chelating agent triethylenetetramine 70 nM (-Cu + Che). The experiment was conducted in 50 mL cultures using plastic flasks, illuminated at 30-60 µmol photons $m^{-2}s^{-1}$E, and shaken at 120 rpm.*

No significant differences were observed in the growth of *Synechococcus elongatus* WT and *ΔcutA::kan* strains in BG11 medium, regardless of the addition of $CuSO_4$. Both strains also grew in the presence of the copper chelator triethylenetetramine, indicating that copper depletion did not significantly impact their viability. These results suggest that *S. elongatus* PCC 7942 compensates for copper limitation by switching from copper-dependent enzymes, such as plastocyanin (involved in photosynthetic electron transport), to iron-dependent alternatives like cytochrome $c_6$. This adaptive mechanism enables the cells to maintain efficient photosynthesis and electron transport under copper-starved conditions, supporting continued growth. Remarkably, the cells fully recovered after the experiment, even while remaining in copper-depleted medium.

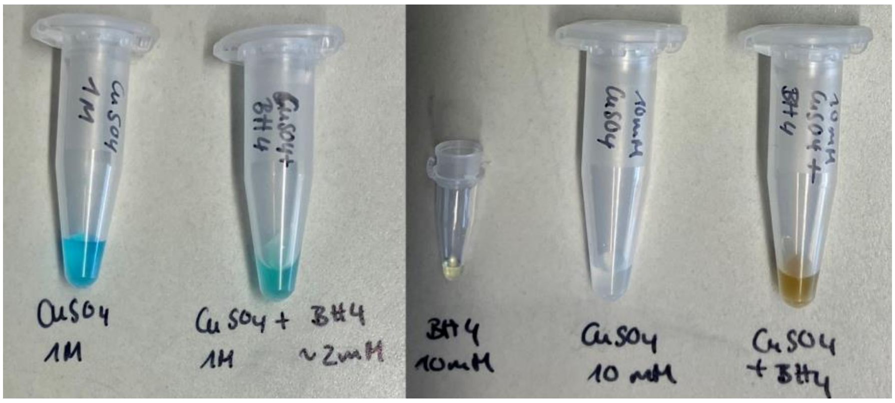

**Supplementary Figure 15: Interaction between $CuSO_4$ and $BH_4$.** Microcentrifuge tubes containing $CuSO_4$ and $BH_4$ are shown, with the $CuSO_4$ solution displaying a characteristic blue color, and the $BH_4$ solution remaining clear. When $CuSO_4$ and $BH_4$ are combined 1:1 in a single tube, the resulting mixture appears brown, indicating a reaction or complex formation between $CuSO_4$ and $BH_4$. This color change suggests a potential interaction between copper ions and $BH_4$.

```
cutA_200bp            1 caaccatccgttcttcatcccgggtactctgggtgttgccctggccgtttgcttcggctt
DQO017_24580176_    192 ----------------------------------------------tttt----------
DQO016_24580169_        ------------------------------------------------------------

                                       cutA_fw
cutA_200bp           61 cgtgctgggtagcttcatgctgtaatgatcaatcgcggggcgttcacg-ccccgctttct
DQO017_24580176_    188 ---------ttgcttcatgctgtaatgatcaatcgcggggcgttcacggccccgctttct
DQO016_24580169_        ------------------------------------------------------------

cutA_200bp          120 ttc-ccgccgactaacatccttcccccgtccgtt-gtatagtgacctctctcttgcggtt
DQO017_24580176_    137 ttttccgccgactaacatccttcccccgtccgtt-gtatagtgacctctctcttgcggtt
DQO016_24580169_      1 -------------------------------gttagtatagtgacctctctcttgcggtt
                          Sequence before cutA
cutA_200bp          178 ccatctgttcttgcgaggtgtttatgcttgatgaaaaaagttcgaataccgcgtctgtcg
DQO017_24580176_     78 ccatctgttcttgcgaggtgttt-------------------------------------
DQO016_24580169_     30 ccatctgttcttgcgaggtgttt-------------------------------------
                                                             cutA
cutA_200bp          238 tggtgctatgtacggcaccagatgaagcgacagcccaggatttagccgccaaagtgctgg
DQO017_24580176_        ------------------------------------------------------------
DQO016_24580169_        ------------------------------------------------------------

cutA_200bp          298 cggaaaaactggcggcctgcgcgaccttgatccccggcgctacctctctctattactggg
DQO017_24580176_        ------------------------------------------------------------
DQO016_24580169_        ------------------------------------------------------------

cutA_200bp          358 aaggtaagctggagcaagaatacgaagtgcagatgattttaaaaactaccgtatctcacc
DQO017_24580176_        ------------------------------------------------------------
DQO016_24580169_        ------------------------------------------------------------

cutA_200bp          418 agcaggcactgctggaatgcctgaagtctcatcatccatatcaaaccccggaacttctgg
DQO017_24580176_        ------------------------------------------------------------
DQO016_24580169_        ------------------------------------------------------------

cutA_200bp          478 ttttacctgttacacacggagacacagattacctctcatggctcaacgcatctttacgct
DQO017_24580176_        ------------------------------------------------------------
DQO016_24580169_        ------------------------------------------------------------
                            Sequence after cutA
cutA_200bp          538 gatcctgctactttgcagcacttccgtttttgccggattattcgacgcgccgggacgttc
DQO017_24580176_     55 --tcctgctactttgcagcacttccgtttttgccggattattcgacgcgccgggacc---
DQO016_24580169_     53 --tcctgctactttgcagcacttccgtttttgccggattattcgacgcgccgggacgttc

cutA_200bp          598 acaatttgtccccgcggatcaagcctttgcttttgattttcagcaaaaccaacatgacct
DQO017_24580176_        ------------------------------------------------------------
DQO016_24580169_    111 acaatttgtccccgcggatcaagcctttgcttttgattttcagcaaaaccaacatgacct
                                        cutA_rev
cutA_200bp          658 taatctgacctggcagatcaaagacggttactacctctaccgtaaacagatccgcattac
DQO017_24580176_        ------------------------------------------------------------
DQO016_24580169_    171 taatctgacctggcagatcaaagacgg------------------------------tac
```

**Supplementary Figure 16: Sequence alignment of *E. coli* ΔcutA after cloning.** Primers flanking the *cutA* sequence were used for sequencing. Alignment of the regions upstream and downstream of *cutA*, but the absence of alignment within the *cutA* sequence, confirms successful seamless cloning of *E. coli* ΔcutA. Forward primer *cutA_fw* (tagcttcatgctgtaatgatcaatcgc) and reverse primer *cutA_rev* (tagaggtagtaaccgtctttgatctgcc) were used for sequencing. The *cutA* sequence along with 200 bp upstream and/or downstream was included for comparison.

### Purification of Proteins

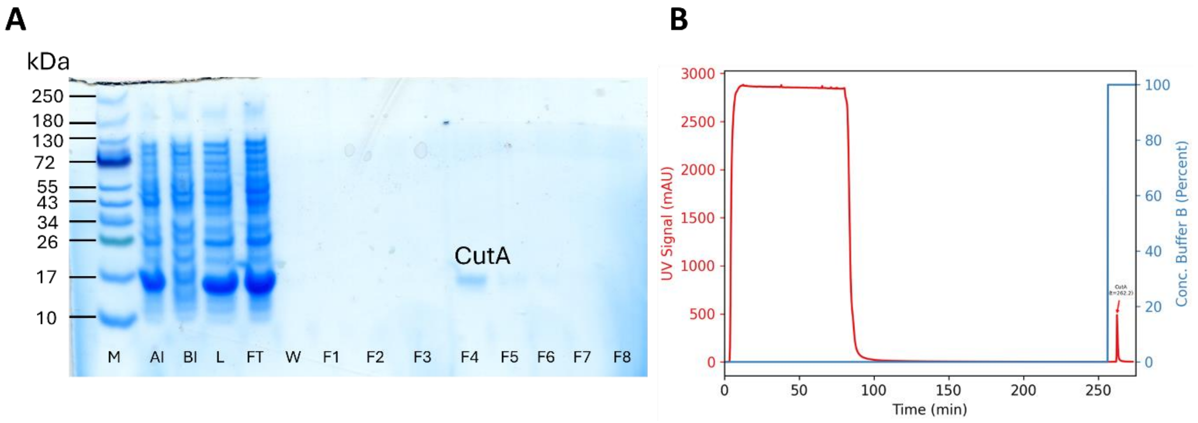

**Supplementary Figure 17: Purification of *E. coli* CutA** *(A) SDS-PAGE analysis of protein samples collected during the purification of E. coli CutA with a Strep-tag. The theoretical molecular weight of the CutA protein, including the Strep-tag, is 13.53 kDa. The samples include: protein marker (M; Color Prestained Protein Standard, Broad Range, 10–250 kDa; NEB P7719S), lysates before induction (BI), after induction (AI), total lysate (L), flow-through (FT), wash fractions (W), and elution fractions (F1–F8). Proteins were resolved on a 14% SDS-PAGE gel. (B) Elution profile from the ÄKTA purifier system showing the UV absorbance at 280 nm and the gradient of Buffer B used for protein elution.*

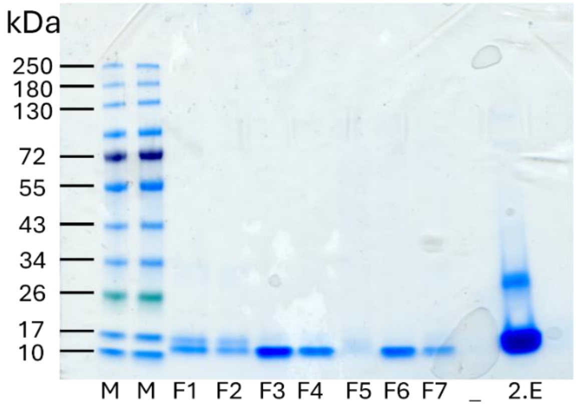

**Supplementary Figure 18: Purification *S. elongatus* CutA** *SDS-PAGE of different collected fractions after purification of E. coli CutA with a Strep-tag. The theoretical mass of the CutA protein, including the Strep-tag, is 13.69 kDa. Samples include protein marker (M; Color Prestained Protein Standard, Broad Range, 10–250 kDa; NEB P7719S), lysates before (BI) and after induction (AI), total lysate (L), flow-through (FT), wash fractions (W), and collected fractions (F1–F7), 2.E = second loading and elution of the column. Proteins were separated using a Q-PAGE™ TGN Precast Gel (Midi, 12 wells, 4–15%).*

### Bacterial strains, Plasmids and Primer

*Table 1: Bacterial Strains Used in This Study*

| Strain | Genotype | Purpose |
|---|---|---|
| *Synechococcus elongatus* PCC 7942 | WT | Characterisation, Pteridine Purification |
| *Synechococcus elongatus* PCC 7942 *ΔcutA::kan*[1] | *ΔcutA::kan* | Characterisation |
| *E. coli* BW225113 | WT | Characterisation |
| *E. coli* BW225113 *ΔcutA (seamless)* | *E.coli* ΔcutA | Characterisation |
| *E. coli* Lemo21(DE3) | *fhuA2 [lon] ompT gal (λ DE3) [dcm] ΔhsdS/* pLemo(Cam$^R$) *λ DE3 = λ sBamHIo ΔEcoRI-B int::(lacI::PlacUV5::T7 gene1) i21 Δnin5* pLemo = pACYC184-*PrhaBAD-lysY* | Protein purification (purchased) |
| *NEB 10-beta Competent E. coli* | *Δ(ara-leu) 7697 araD139 fhuAΔlacX74 galK16 galE15 e14-Φ80dlacZΔM15 recA1 relA1 endA1 nupGrpsL(StrR) rphspoT1 Δ(mrr-hsdRMS-mcrBC)* | Plasmid propagation for molecular cloning of larger plasmids or difficult constructs (purchased) |
| *E. coli* TOP10 *Competent Cells* | F$^-$ *mcrA Δ(mrr-hsdRMS-mcrBC) φ80lacZΔM15 lacX74 recA1 araD139 Δ(ara-leu)7697 galU galK rpsL (Str$^r$) endA1 nupG* | Plasmid propagation for molecular cloning of smaller plasmids (purchased) |

*Table 2: Plasmids Used in This Study*

| Plasmid | Resistance | Purpose |
|---|---|---|
| **pSLTS** [2] | Amp (100) | Molecular cloning *ΔcutA, carries a selection marker, I-SceI cut site, and a transcription terminator (TT)* |
| **pT2SK**[2] | Ampicillin (100) and Kanamycin (50) | Molecular cloning *ΔcutA,* carries lambda-red recombinase + I-SceI for homologous recombination |
| pASK[*E.coli_cutA*] | Ampicillin (100 µg/mL) | Protein purification |
| pASK[*S. elongatus_cutA*] | Ampicillin (100 µg/mL) | Protein purification |

*Table 3: Primers Used in This Study*

| Primer | Sequence (5`→3´) | Purpose |
|---|---|---|
| P1_*cutA*_fw[1] | TAGCTTCATGCTGTAATGATCAATCGCG | Sequencing of *E. coli cutA* |
| P2_*cutA*_rev[1] | TAGAGGTAGTAACCGTCTTTGATCTGCC | Sequencing of *E. coli cutA* |
| pHA.seq.F | TATCAGGGTTATTGTCTCATGAGCG | Molecular cloning *ΔcutA** |
| pHA.seq.R | ACTTGAGCGTCGATTTTTGTGATGC | Molecular cloning *ΔcutA* |
| pKDTS-F | TAGGCGCAATCACTTTCGTCTACTC | Molecular cloning *ΔcutA* |
| pKDTS-R | TTGAGTGACATGCAAAGTAAGTATGATCTC | Molecular cloning *ΔcutA* |
| pHAFor | CGCAGGAAAGAACATGTG | Molecular cloning *ΔcutA* |
| pHARev | AAGGGCCTCGTGATACG | Molecular cloning *ΔcutA* |

| | | |
|---|---|---|
| MF | ATCTCAAGAGTGGCAGC | Molecular cloning *ΔcutA* |
| MR | TTACGCCCCGCCCTGC | Molecular cloning *ΔcutA* |
| 5`- mut cassette _fw | AGGCGTATCACGAGGCCCTTATGATCAATCGC GGGGCGTTCAC | Molecular cloning *ΔcutA* *5´mutation fragment* |
| 5`- mut. Cassette rev | ACCGCTGCCACTCTTGAGATTGCAAAGTAGCA GGAAAACACCTCGCAAGAACAGATGGAACCG | Molecular cloning *ΔcutA* *5'mutation fragment* |
| 3´mut. Cassette fw | GCAGGGCGGGGCGTAATCTTGCGAGGTGTTT TCCTGCTACTTTGCAGCACTTCCGTTTTTG | Molecular cloning *ΔcutA* *3'mutation fragment* |
| 3`mut. Cassette rev | CTCACATGTTCTTTCCTGCGCATGTTGGTTTTG CTGAAAATC | Molecular cloning *ΔcutA* *3'mutation fragment* |
| *E. coli*_cutA_Gibson_fw | GTGAAATGAATAGTTCGACAAAAATCTAGATA ACGAGGGCAAAAA**ATGCTTGATGAAAAAAG** | Molecular cloning pASK [*E. coli_cutA1*] for protein purification |
| *E. coli*_pASK_cutA_Gibson_rev | CTTATTATTTTTCGAACTGCGGGTGGCTCCAAG CGCT**GCGTAAAGATGCGTTGAGCCATG** | Molecular cloning pASK [*E. coli_cutA1*] for protein purification |
| pASK_IBA3_Fw | GAGTTATTTTACCACTCCCT | Sequencing of pASK plasmids |
| pASK_IBA3_Rv | ACGCAGTAGCGGTAAAC | Sequencing of pASK plasmids |

1. Selim, K. A. *et al.* Functional and structural characterization of PII-like protein CutA does not support involvement in heavy metal tolerance and hints at a small-molecule carrying/signaling role. *FEBS J.* **288**, 1142–1162 (2021).
2. Kim, J., Webb, A. M., Kershner, J. P., Blaskowski, S. & Copley, S. D. A versatile and highly efficient method for scarless genome editing in Escherichia coli and Salmonella enterica. *BMC Biotechnol.* **14**, 84 (2014).
